# Th1 cells alter the inflammatory signature of IL-6 by channeling STAT transcription factors to *Alu*-like retroelements

**DOI:** 10.1101/2022.07.18.499157

**Authors:** David Millrine, Ana Cardus Figueras, Javier Uceda Fernandez, Robert Andrews, Barbara Szomolay, Benjamin C Cossins, Christopher M. Rice, Jasmine Li, Victoria J Tyrrell, Louise McLeod, Peter Holmans, Valerie B O’Donnell, Philip R Taylor, Stephen J. Turner, Brendan J. Jenkins, Gareth W Jones, Nicholas Topley, Nigel M Williams, Simon A Jones

## Abstract

Cytokines that signal via STAT1 and STAT3 transcription factors instruct decisions affecting tissue homeostasis, anti-microbial host defense, and inflammation-induced tissue injury. To understand the coordination of these activities, we applied RNA-seq, ChIP-seq, and ATAC-seq to identify the transcriptional output of STAT1 and STAT3 in peritoneal tissues during acute resolving inflammation and inflammation primed to drive fibrosis. Bioinformatics focussed on the transcriptional signature of the immuno-modulatory cytokine IL-6 in both settings and examined how pro-fibrotic IFNγ-secreting CD4^+^ T-cells altered the interpretation of STAT1 and STAT3 cytokine cues. In resolving inflammation, STAT1 and STAT3 cooperated to drive stromal gene expression affecting anti-microbial immunity and tissue homeostasis. The introduction of IFNγ-secreting CD4^+^ T-cells altered this transcriptional program and channeled STAT1 and STAT3 to a previously latent GAS motif in *Alu*-like elements. STAT1 and STAT3 binding to this conserved sequence revealed evidence of reciprocal cross-regulation and gene signatures relevant to pathophysiology. Thus, we propose that effector T-cells re-tune the transcriptional output of IL-6 by shaping a regulatory interplay between STAT1 and STAT3 in inflammation.

## Introduction

Cytokines are essential for development, tissue homeostasis, and the regulation of inflammation (Dinarello, 2007). The intracellular signaling pathways controlling these activities are intrinsically linked, and aberrant host defense compromises tissue integrity and physiological function. Patients treated with anti-cytokine therapies often provide evidence of this relationship (Choy *et al*, 2013). For example, interleukin (IL)-6 inhibition is less effective in diseases where IL-6 maintains tissue homeostasis and barrier immunity (Andrews *et al*, 2018; Harbour *et al*, 2020; Hunter & Jones, 2015; Jones & Jenkins, 2018; Jones *et al*, 2011; Navarini *et al*, 2011; Schett *et al*, 2013; Taniguchi *et al*, 2015). How IL-6 operating through a single receptor signaling cassette coordinates the maintenance of tissue physiology and the transition to pathophysiology is unknown.

Interleukin-6 regulates cellular responses *via* receptor activation of Janus kinases (Jak) and members of the Signal Transducer and Activator of Transcription (STAT) family (Hunter & Jones, 2015; Jones & Jenkins, 2018). While IL-6 employs other signaling intermediates, IL-6 instructs cell decisions primarily through STAT1 and STAT3 transcription factors (Costa-Pereira *et al*, 2002; Hirahara *et al*, 2015; Jones *et al*, 2013; Peters *et al*, 2015; Twohig, 2019; Villarino *et al*, 2017; Wilmes *et al*, 2021). These proteins share a complex regulatory interplay, and gene ablation studies show that STAT1 and STAT3 often counteract each other or engage shared enhancers (Avalle *et al*, 2012; Hunter & Jones, 2015; Martinez-Fabregas *et al*, 2020; Vahedi *et al*, 2015). These interactions re-tune the interpretation of cytokine cues, instructing alternate patterns of gene regulation (Fielding *et al*, 2014; Grohmann *et al*, 2018; Hong *et al*, 2002; Jones *et al*, 2015; Jones et al., 2013; Lin & Leonard, 2019; Martinez-Fabregas et al., 2020; Twohig, 2019; Wiede *et al*, 2019).

Resident tissue cells respond to immune challenge by steering decision-making processes affecting disease outcome (Dinarello, 2007; Gomes & Teichmann, 2020; Jones, 2005; Krausgruber *et al*, 2020). These activities rely on cytokine networks that promote communication between stromal tissue and infiltrating leukocytes (Jones, 2005). In bacterial peritonitis, IL-6 controls anti-microbial immunity and the resolution of inflammation by steering the transition from innate to adaptive immunity (Catar *et al*, 2017; Cho *et al*, 2014; Hurst *et al*, 2001; Jones *et al*, 2010; McLoughlin *et al*, 2004; McLoughlin *et al*, 2005). This process requires IL-6R shedding from infiltrating neutrophils, which promotes IL-6 trans-signaling and STAT3-driven outcomes in resident stromal cells (Fielding *et al*, 2008; Hurst et al., 2001; Jones *et al*, 1999; McLoughlin et al., 2004; McLoughlin *et al*, 2003). Repeated immune activation alters this response, with IL-6 supporting an expansion of pro-fibrotic IFNγ-secreting Th1 cells, which initiate tissue injury through activation of stromal STAT1 signaling (Fielding et al., 2014). To understand how the effector properties of Th1 cells impact the contribution of STAT1 and STAT3 in acute resolving peritonitis, we have applied next-generation sequencing methods to examine the stromal response to inflammation. Our analysis shows that Th1 cells alter the transcriptional output of IL-6 by re-directing STAT transcription factors to a GAS-like motif in *Alu*-like retroelements. These results offer an explanation of how effector lymphocytes shape the stromal response to inflammation by modifying the interpretation of cytokine cues.

## Results

### Effector Th1 cells shape stromal responses to peritonitis

To understand how pro-fibrotic Th1 cells shape the stromal cell response to acute peritonitis, we challenged mice with an intraperitoneal (i.p.) administration of cell-free supernatant from *Staphylococcus epidermidis* (SES) (Fielding et al., 2014; Hurst et al., 2001; Jenkins *et al*, 2005). Mice received SES alone or co-administered with CD4^+^ T-cells expanded *ex vivo* into Th1 cells (normalized to 10^6^ IFNγ-secreting T-cells) (Fielding et al., 2014). To control for transcriptional changes induced by Th1 cells (SES+Th1), a separate group of mice received SES and an equivalent number of naïve CD4^+^ T-cells (Fig-1). At 3 and 6 hours post-stimulation, the peritoneum was harvested and prepared for RNA-seq. K-means clustering was restricted to transcripts differentially regulated (Log2FC>1.75, P_adj_<0.05) under each condition (Fig-1A). Applying statistical tools (silhouette, gap statistics, and elbow), we identified four clusters of gene regulation in mice receiving SES alone or SES+Th1 (Fig-1B). We identified 821 differentially regulated transcripts whose expression altered in (at least) one time point. SES regulated a total of 225 genes, with the activities of Th1 cells affecting another 673 genes. These included genes affecting vascularization, epithelial morphogenesis, and hyperplasia (e.g., *Angpt4, Egr3, Igfbp2, Ntf3, Tbxa2r*). Others displayed involvement in innate sensing pathways (e.g., *Ifitm1, Marco, Myd88*) and leukocyte infiltration (e.g., *Ccl3, Ccl5, Ccl2, Cxcl1, Cxcl10, Icam1, Vcam1*) (Fig-1C). The presence of Th1 cells altered gene expression in each cluster, with molecular pathway analysis highlighting an enrichment of transcripts attributed to STAT1 and interferon signaling (Fig-2A & 2B; Supplemental Figure-1). To substantiate these findings, we used the Upstream Regulator tool in Ingenuity Pathway Analysis to predict transcriptional mechanisms accounting for changes in gene expression in each dataset. Consistent with the SES activation of TLR2 (Colmont *et al*, 2011), this algorithm identified genes controlled by the NF-κB pathway (Fig-2C). It also identified genes linked with STAT1, STAT3, and several IRF transcription factors (Fig-2C). Transcripts affiliated with STAT1 and STAT3 showed considerable enrichment in SES+Th1 treated mice, suggesting a potential link between IL-6 and IFNγ signaling in steering SES-induced outcomes (Fielding et al., 2014; Hurst et al., 2001; McLoughlin et al., 2003; Robson *et al*, 2001).

**Figure 1.**
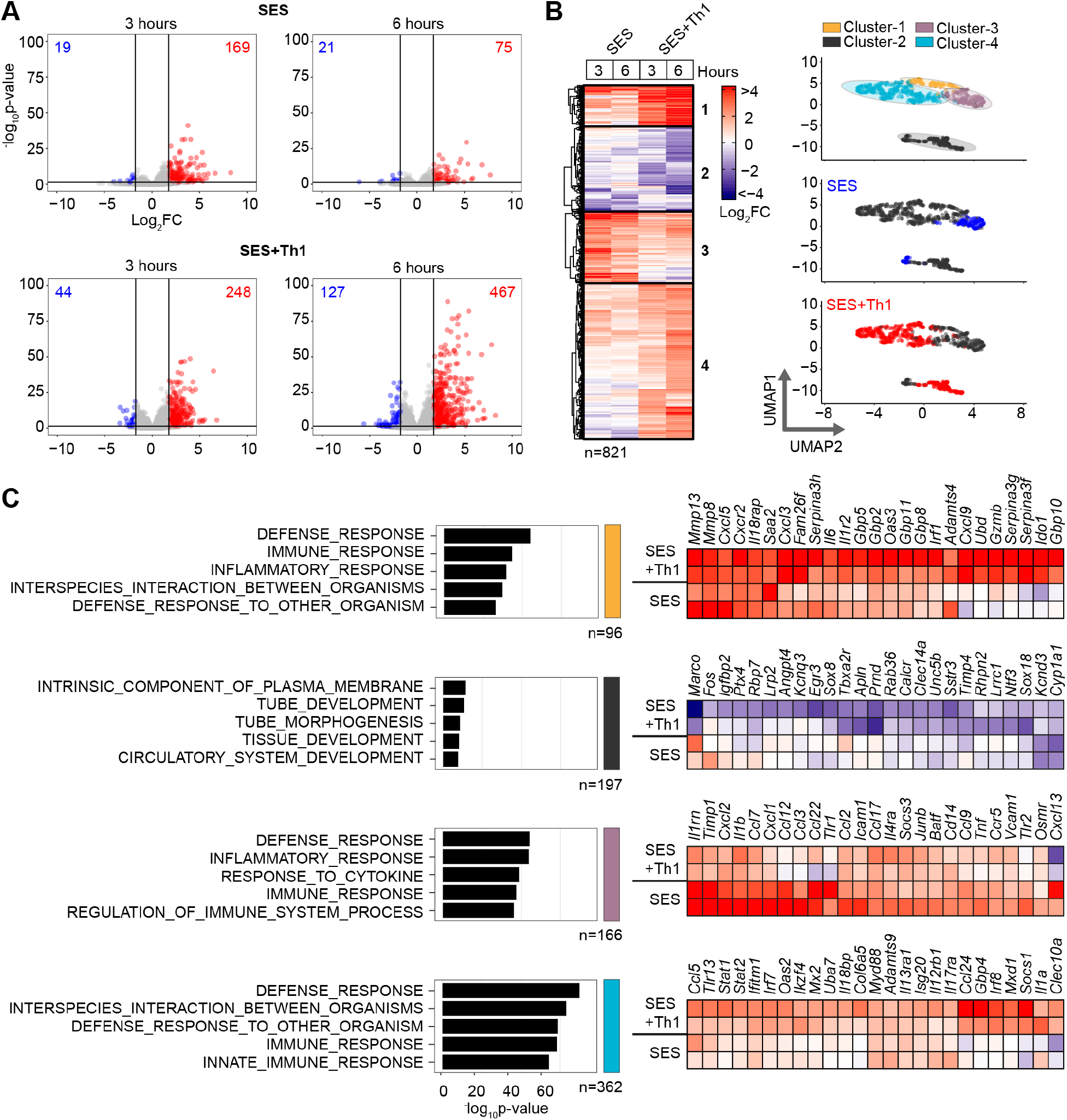
Th1 cells augment the stromal response to acute inflammation. **(A)** RNA-seq was performed on stromal tissues extracted from SES challenged mice. Volcano plots show differential expression analysis (LIMMA) and statistical thresholding (p_adj_<0.05, Log_2_FC>1.75) of datasets at 3 and 6 hours post treatment. Mice received (i.p.) SES alone (SES) or SES in combination with Th1 cells (SES+Th1). Each test condition was compared to untreated mice (SES), or mice receiving SES and naïve CD4+ T-cells. **(B)** K-means clustering (n=4) of data shown in Panel-A (heatmap). UMAP visualizations show the distribution of gene clusters and their link to SES (blue) or SES+Th1 (red) datasets. **(C)** Gene Ontology (msigdb) shows the top-5 biological processes for each cluster (left). Each cluster descriptor is coded to match UMAP clustering colors in Panel-B. Representative examples of genes in each cluster are presented as heatmaps (right).

**Figure 2.**
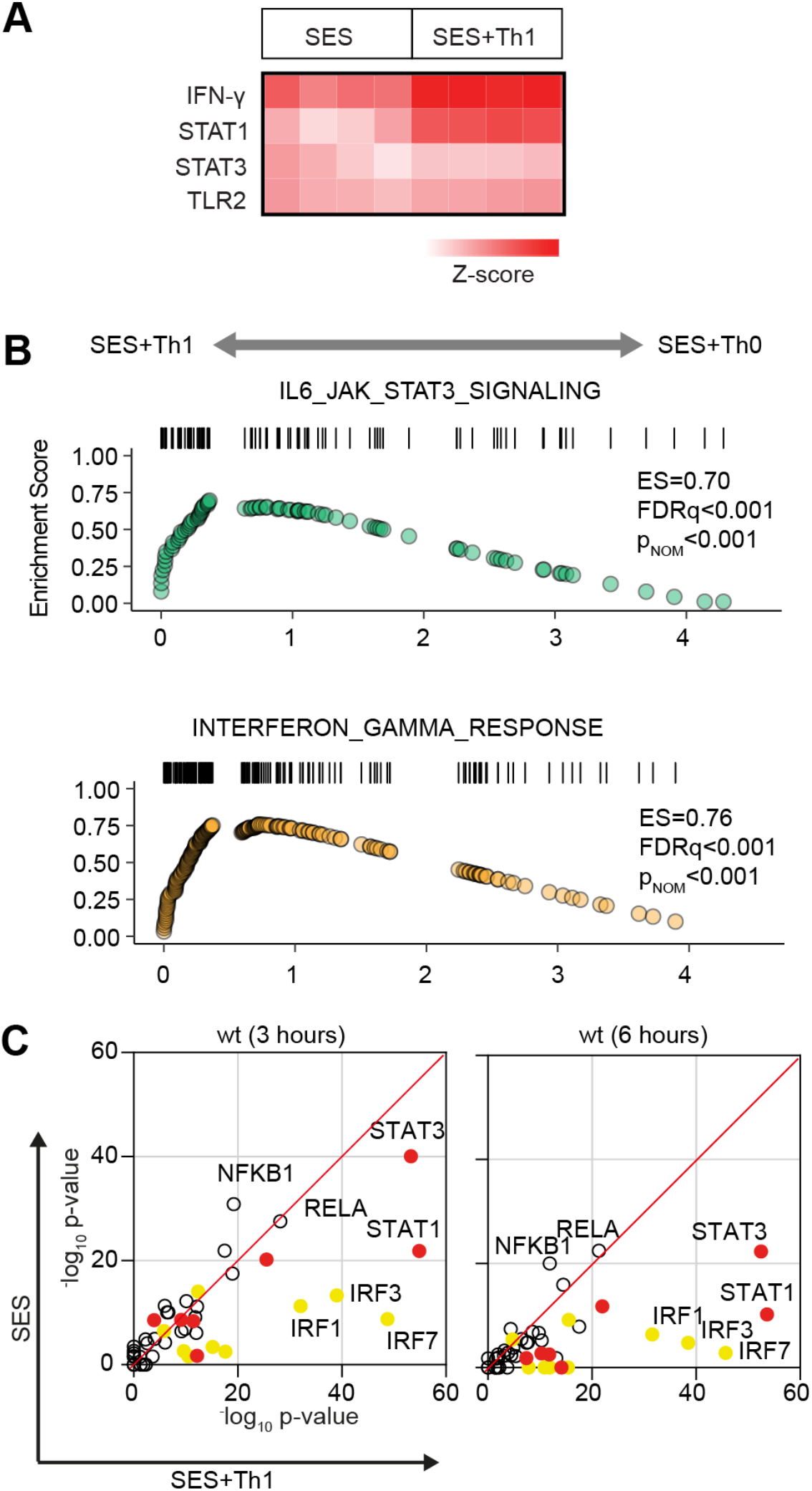
Regulatory signatures of cytokine signalling in acute inflammation. **(A)** Ingenuity Pathway Analysis of biological processes involved in SES-induced inflammation. **(B)** Gene set enrichment analysis shows the impact of Th1 cells on Jak-STAT cytokine signalling and interferon regulated outcomes. A summary of the complete GSEA analysis is provided in ***Supplemental Figure-1***. **(C)** Ingenuity Pathway Analysis of upstream regulators. Predicted p-value (Fisher’s exact test) identifies gene signatures characteristic of specific transcription factors linked to TLR (NF-κB, Rel-A and shown), interferon (Yellow; IRF family members are shown) and Jak-STAT cytokine signaling (Red; STAT1 and STAT3 are displayed).

### Th1 cells modify the properties of IL-6

To explore the link between IL-6 and IFNγ in SES-induced peritonitis, we used immunodetection methods to quantify changes in their production as a reponse to SES challenge (Supplemental Figure-2A). Peritoneal IFNγ remained below the limit of detection in mice treated with SES alone. However, IFNγ levels were substantially enhanced by the introduction of Th1 cells (Supplemental Figure-2A). The presence of Th1 cells also increased IL-6 bioavailability by promoting 3-4-fold increases in IL-6 and sIL-6R (Supplemental Figure-2A). To establish if Th1 cells alter the transcriptional output of IL-6, we performed RNA-seq on peritoneal tissues from SES challenged wild type (*wt*) and *Il6*^−/-^ mice lacking an ability to signal via classical IL-6R signaling and IL-6 trans-signaling. K-means clustering identified 241 significantly altered transcripts (Log2FC>1.75, P_adj_<0.05) impacted by the loss of IL-6 (Fig-3A & 3B, Supplemental Figure-2B-D). These included gene required for tissue homeostasis (e.g., *Atoh1, Cldn5, Fgfbp1, Npnt, Oxtr, Pdx1, Pgr, Stab2*), host defense (e.g., *Trim52, C7*), leukocyte recruitment (e.g., *Ccl8, Ccl17, Ccl22, Ccl24*), and adhesion (e.g., *Selp, Adgrb2*) (Fig-3C). To identify the peritoneal stromal cells responding to IL-6 in SES-induced inflammation, we compared our results against a single cell RNA-seq dataset (GEO GSM4053741) from mouse omental CD45^−^CD41^−^Ter119^−^CD31^−^PDPN^+/−^ cells (Jackson-Jones *et al*, 2020). This analysis defined roles for peritoneal fibroblasts (including *Ccl11^+^ Pdgfra^+^ and Matn2^+^ Pdgfra^+^* subsets) and mesothelial cells (including *Ifit*^+^ and *Cxcl13*^+^ mesothelial cells) in shaping IL-6 responses (Supplemental Figure-2E).

**Figure 3.**
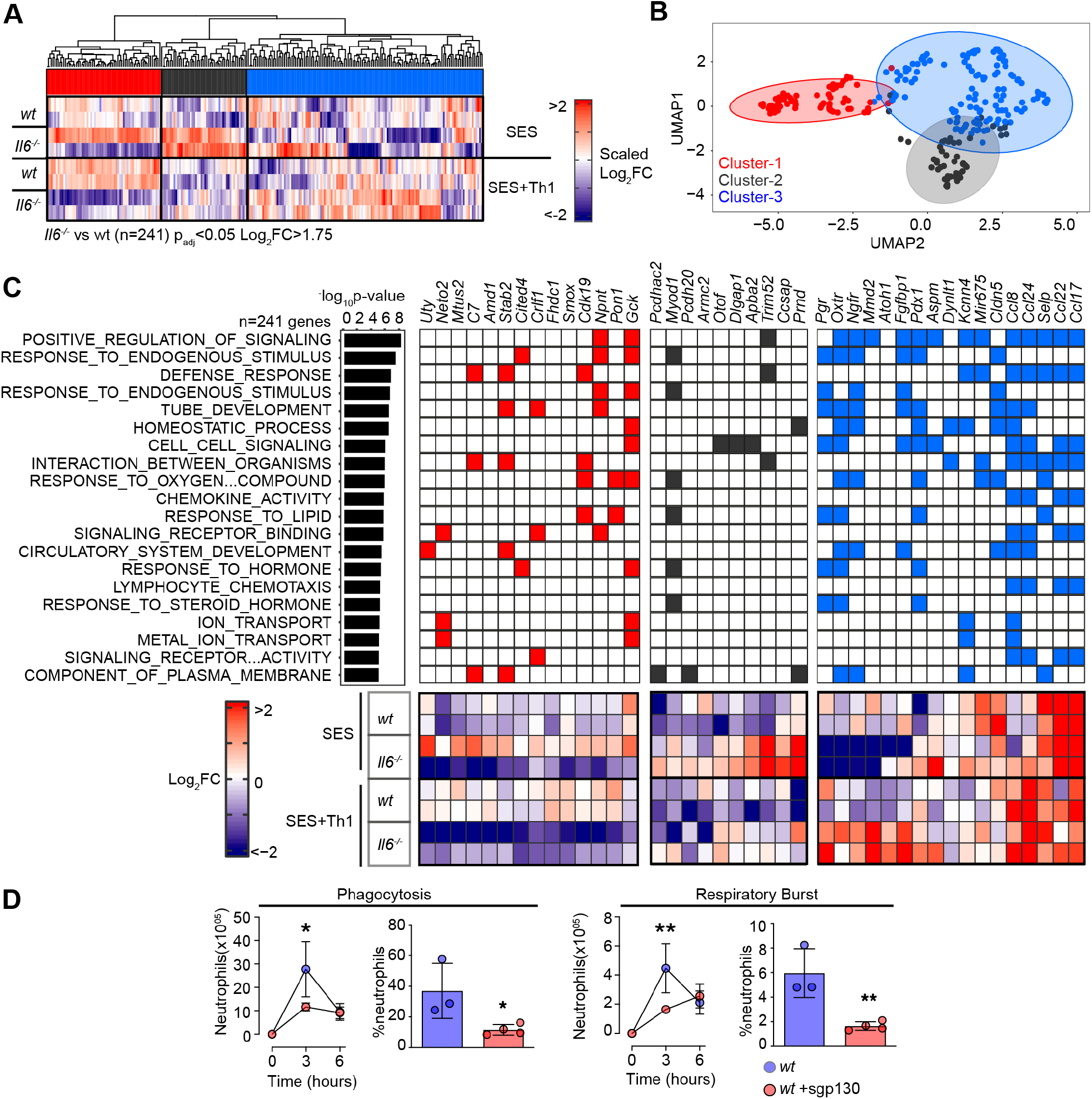
Th1 cells shape the transcriptional output of IL-6. **(A)** Heatmap of IL-6 regulated transcripts identified by differential expression analysis (LIMMA) of *wt* versus *Il6−/−* mice. Data is arranged by K-means clustering (n=3) of significantly regulated transcripts (Cluster-1; red, Cluster-2; grey, Cluster-3; blue). **(B)** UMAP visualization of all IL-6 regulated transcripts (p_adj_<0.05, Log_2_FC>1.75). **(C)** Alignment of representative transcripts from each cluster against the top-20 biological processes identified by gene ontology enrichment analysis of 241 IL-6 regulated transcripts (msigdb). **(D)** Flow cytometric analysis of infiltrating neutrophil effector function (see ***Supplemental Figure-2***). Peritonitis was induced in WT mice by administration of fluorescently labelled Staphylococcus epidermidis (5×10^8^ CFU) in the presence of sgp130Fc (250ng/mouse). Changes in phagocytosis and respiratory burst are shown (mean ± SEM, n=4; **p<0.01, *p<0.05).

Our initial bioinformatic predictions suggested that IL-6 controls stromal responses affecting innate immunity. To verify this connection, we developed a flow cytometric method to compare the effector properties of circulating and infiltrating Ly-6B^hi^ Ly-6G^hi^ neutrophils from *wt* and *Il6*^−/-^ mice. Circulating neutrophils were loaded *ex vivo* with the peroxidase substrate 3’(p-aminophenyl) fluorescein (APF) and exposed to opsonized *Staphylococcus epidermidis* labeled with DDAO-far-red. These reporter dyes were used to track neutrophil respiratory burst and phagocytosis capabilities. Circulating Ly-6B^hi^ Ly-6G^hi^ neutrophils from wt and *Il6*^−/-^ mice showed no differences in effector functions (Supplemental Figure-2F). However, infiltrating neutrophils from *Il6*^−/-^ mice treated (i.p.) with 5×10^8^ cfu fluorescently labeled *Staphylococcus epidermidis* displayed impaired neutrophil function (Fig-3D, Supplemental Figure-2G). We further confirmed these findings by visualizing fluorescent bacteria in neutrophils using imaging flow cytometry (Supplemental Figure-2H). Since IL-6 requires sIL-6R to regulate stromal responses within the peritoneal cavity, we conducted an identical experiment in *wt* mice treated with the IL-6 trans-signaling antagonist soluble gp130 (sgp130). Treatment with sgp130 significantly reduced the effector properties of infiltrating neutrophils in infected mice (Fig-3D). This defect was reversed by reconstituting IL-6 signaling (via the administration [i.p.] of a chimeric IL-6-sIL-6R fusion protein) in *Il6*^−/-^ mice (Fig-3D, Supplemental Figure-2H). Thus, IL-6 governs neutrophil responses to local infection.

### Th1 cells alter STAT transcription factor activity

STAT1 and STAT3 transcription factors become rapidly activated following SES challenge, with maximal activation coinciding with the 3 and 6 hours chosen for our RNA-seq analysis (Fielding et al., 2014; Fielding et al., 2008; McLoughlin et al., 2005) (Supplemental Figure-3A). STAT1 activities often shape the transcriptional output of IL-6 and STAT3 (Costa-Pereira et al., 2002; Twohig, 2019; Vahedi *et al*, 2012; Villarino et al., 2017). To test if this relationship is seen in response to SES, we extracted peritoneal tissues from *gp130*^Y757F:Y757F^ mice challenged with SES. These animals possess a single tyrosine-to-phenylalanine substitution in the cytoplasmic domain of gp130 that prevents the negative regulation of STAT1 and STAT3 following cytokine activation (Jenkins et al., 2005; McLoughlin et al., 2005). Immunoblot for tyrosine-phosphorylated STAT1 (pY-STAT1) and STAT3 (pY-STAT3) showed that SES triggers a prolonged STAT transcription factor activation in these mice (Supplemental Figure-3A). Here, partial Stat3 ablation in *gp130*^Y757F:Y757F^ mice (*gp130*^Y757F:Y757F^:Stat3^+/−^) extended the duration of pY-STAT1 activity in response to SES challenge (Supplemental Figure-3A). Thus STAT1 and STAT3 activities are interlinked in SES inflammation and may explain how Th1 cells impact the transcriptional output of IL-6. We, therefore, applied chromatin immunoprecipitation-sequencing (ChIP-seq) to investigate how STAT1 and STAT3 transcription factors engage the genome following SES challenge (Fig-4, Supplemental Figure-3B). Our analysis identified sequencing peaks displaying a four-fold enrichment above input (P<0.0001; false discovery rate 0.05). These mapped to transcriptional start sites (TSS), exons, introns, and intergenic regions (Supplemental Figure-3C). Motif enrichment analysis (MEME-ChIP) confirmed the specificity of these interactions and identified consensus DNA binding motifs for STAT1 or STAT3 beneath the mapped sequencing peaks (Supplemental Figure-3B).

**Figure 4.**
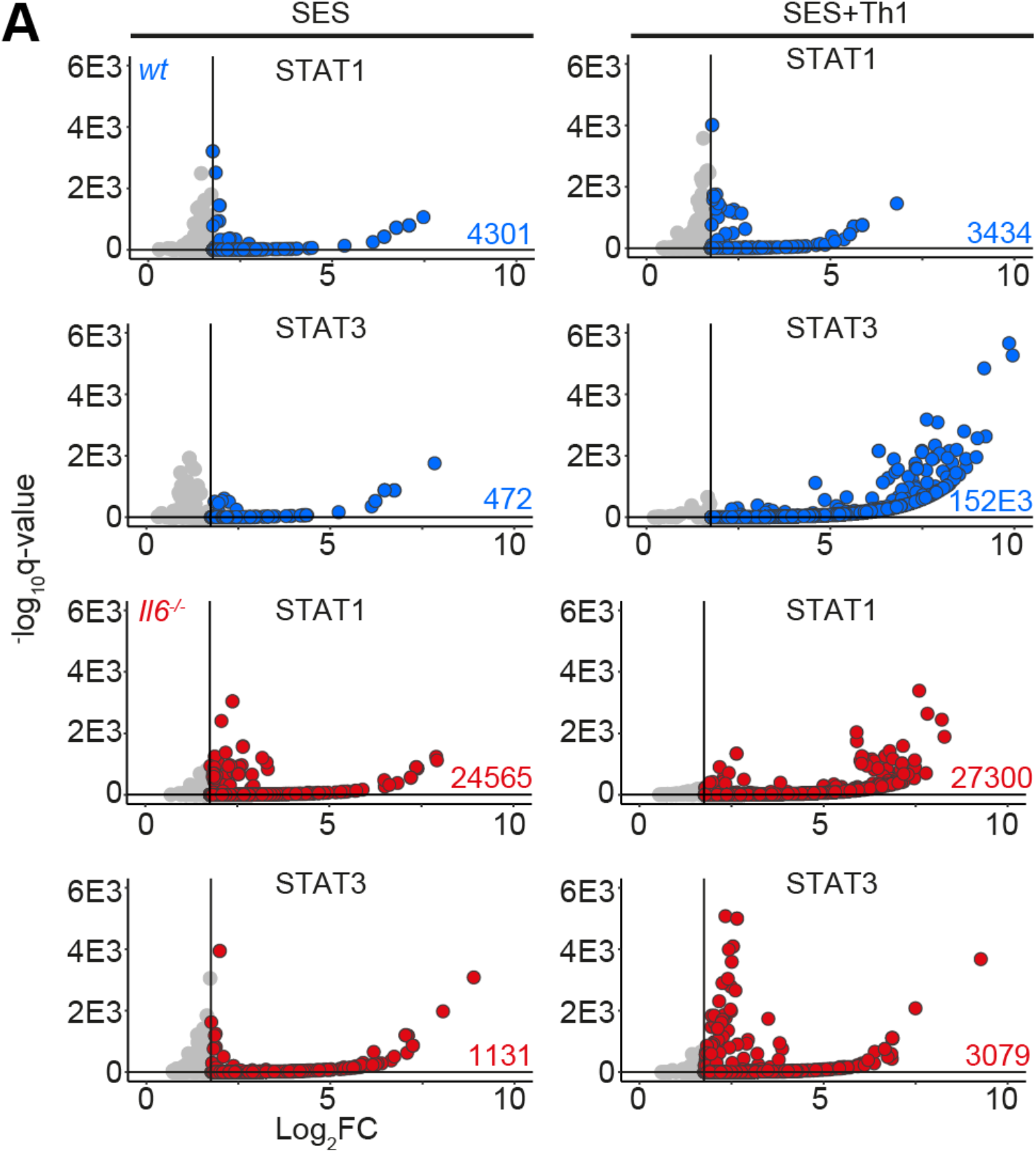
ChIP-seq analysis of STAT1 and STAT3 involvement in SES-induced inflammation. Genomic DNA from the peritoneal membrane of mice challenged with SES alone or SES+Th1 was extracted at 3 hours. Peak calling and downstream processing are described in *Materials and Methods*. Volcano plots summarize ChIP-seq profiling. Each dot represents a peak (Grey = q>0.05 and/or log_2_FC>1.75). Peaks below the significance (q<0.05) and above the log_2_FC (>1.75) cutoff values are highlighted in blue (*wt*) and red (*Il6^−/−^*).

The presence of Th1 cells noticeably altered the genomic localization of STAT transcription factors following SES stimulation (Fig-4A). Following treatment with SES alone, STAT1 and STAT3 worked in close partnership and often bound loci in nearby proximity. Th1 cells increased the number of sequencing peaks identified by ChIP-seq for STAT1 and STAT3 (Supplemental Figure-3C). This increase in STAT binding was consistent with the enhanced gene regulation associated with Th1 cell involvement (Fig-1 & Fig-3). Comparing STAT transcription factor binding in SES+Th1 datasets from wt and *Il6*^−/-^ mice, we also identified genomic loci displaying evidence of STAT1 and STAT3 cross-regulation (Fig-5). Here, STAT transcription factors often bound similar genomic coordinates at downstream sequences distal of TSS. In DNA samples from wt mice, these sites were STAT3 occupied. However, in *Il6* deficiency, these same sites showed increases in STAT1 binding. Promoters displaying this form of cross-regulation included genes involved in cytoskeletal organization (e.g., *Mtus2, Actb, Rhpn2, Wasf1*), metabolism (e.g., *Angptl4, Mtor, Neu2, Pgm1*), and tissue remodeling (e.g., *Timp1, Col2a1, Vegf*) (Fig-5A). This switch from STAT3 to STAT1 coincided with transcriptional changes marked by the induction or suppression of gene expression under Il6 deficiency (Fig-5B). These included the suppression of genes affecting cellular differentiation (e.g., *Dnaic1, Eif2s3y*) and increases in genes linked with matrix protein biosynthesis (e.g., *Acan, Npnt, Col2a1*). Thus, STAT1 and STAT3 coordinate differential impacts on the control of tissue homeostasis.

**Figure 5.**
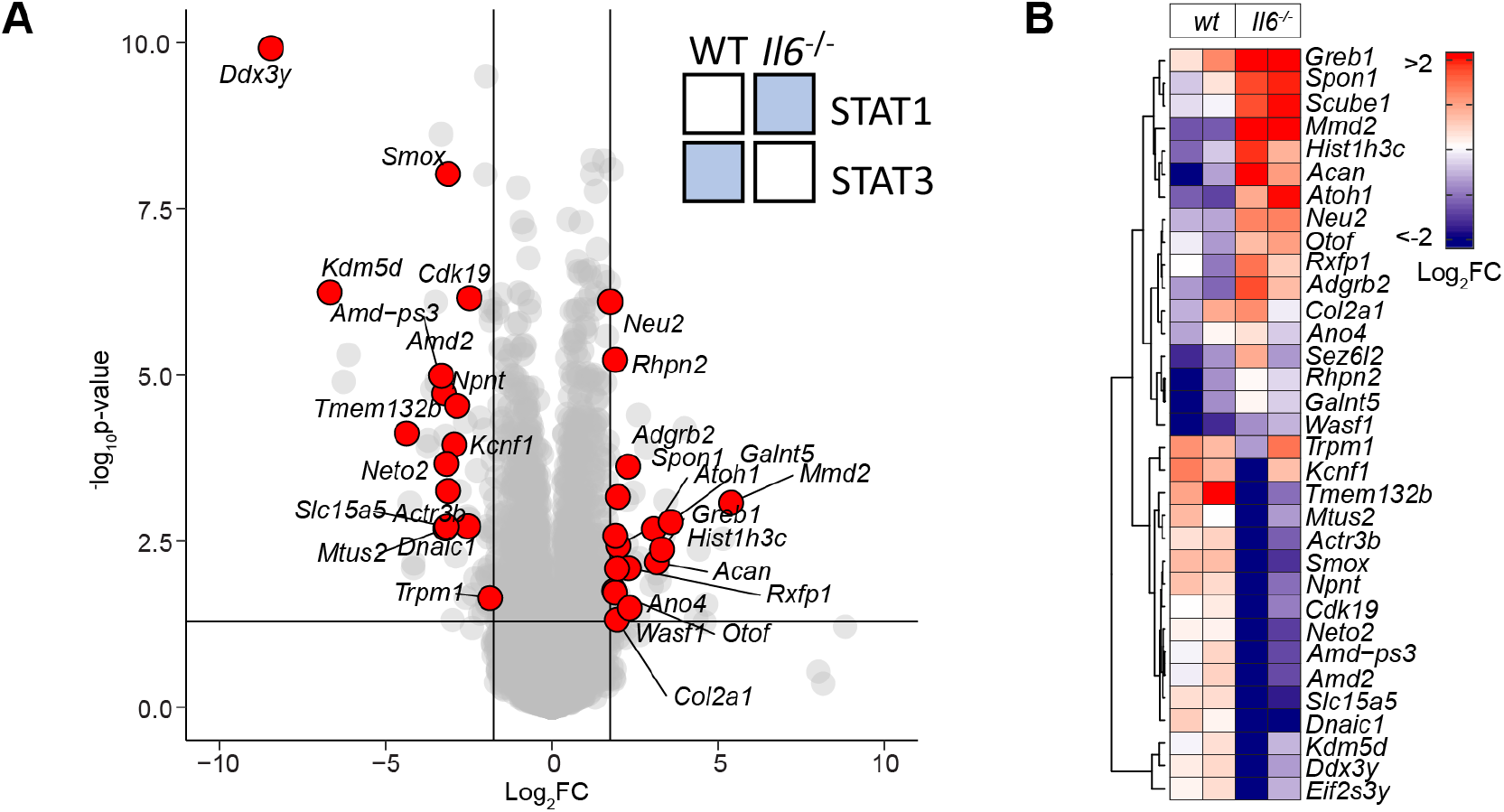
STAT1 and STAT3 interplay shapes gene regulation. **(A)** Volcano plot of RNA-seq data showing differentially regulated gene expression (*Il6*^−/-^ vs *wt*; p_adj_<0.05, Log_2_FC>1.75) in SES+Th1 treated *Il6^−/−^* mice (3-hour post administration). Differential gene regulation is shown for representative genes displaying reciprocal STAT1 and STAT3 binding in ChIP-seq datasets from *wt* and *Il6*^−/−^ mice (summarized in inset). **(B)** Euclidean clustering of the 33 genes depicted in Panel-A.

To substantiate the significance of genomic interactions, we used Assay for Transposase-Accessible Chromatin-sequencing (ATAC-seq) to confirm links between STAT transcription factor binding and chromatin accessibility in SES+Th1 treated mice (Fig-6). The heatmap profiles show chromatin accessibility across the entire genome for wt and *Il6*^−/-^ mice (Fig-6A). Adopting a differential binding analysis (*Diffbind*) of ATAC-seq datasets, we identified open chromatin regions linked with IL-6 bioactivity. These sites showed enrichment at TSS (Fig-6B). Access to these sites was partially restricted by the absence of *Il6*, suggesting that IL-6 promotes chromatin remodeling in inflammation (Fig-6C). Motif enrichment analysis of the DNA sequences aligned to these peaks identified consensus sites consistent with the computational predictions shown in Figure 2C. These included STAT and IRF transcription factors and others (e.g., KLF15, NRF1, HOXOA13, and several zinc-binding factors) contributing to tissue homeostasis and epigenetic control (Fig-6D).

**Figure 6.**
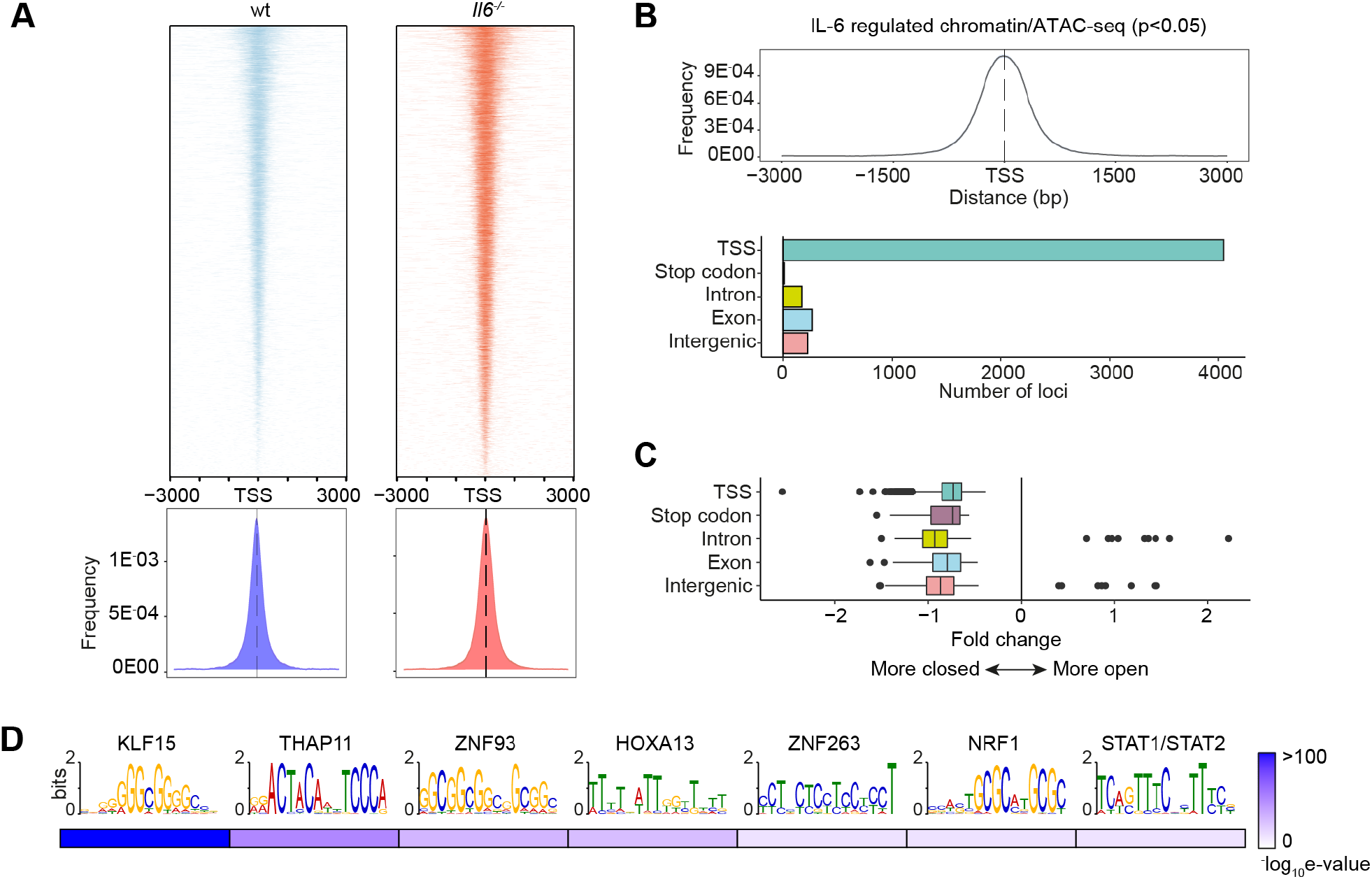
Mapping of chromatin accessibility by ATAC-seq. **(A)** Heatmap visualization of ATAC-seq profiling of peritoneal extracts from mice treated with SES and Th1 cells (Top). The peak count frequency of sequence reads associated with transcription start sites (TSS) is shown for *wt* and *Il6*^−/−^ mice (Bottom). **(B)** Histogram shows chromatin accessibility at TSS linked with IL-6 regulated genes (Top). Graph shows the genomic distribution of IL-6 regulated loci (Bottom). **(C)** Fold change (*Il6*^−/−^ vs *wt*) in ATAC-seq reads at indicated genomic features. **(D)** Motif enrichment analysis (MEME-ChIP) of genomic regions identified in differential binding analysis of ATAC-seq datasets. Annotations identify putative transcription factor motifs associated with sequencing peaks.

### STAT transcription factors engage Alu-like retroelements

The proximity of STAT1 and STAT3 binding to consensus sequences for other transcription factors suggested links to regulatory regions such as super-enhancers. We, therefore, mapped the genomic localization of P300 in peritoneal tissue extracts from mice challenged with SES+Th1 cells (Fig-7A). This histone acetyltransferase controls chromatin remodeling and often localizes active or poised enhancers, where P300 functions as a scaffolding factor and co-regulator of transcription factor activity (Hutchins *et al*, 2013; Villarino et al., 2017). ChIP-seq analysis for P300 identified sequencing peaks sharing STAT transcription factor binding (Supplemental Figure-3B). However, P300 mapping represented a small proportion (<11%) of the total sequencing peaks identified by STAT1 and STAT3 ChIP-seq (Fig-7A). Although 60% of the P300 loci showed a switch in STAT transcription factor binding under *Il6* deficiency, the transition from STAT3 to STAT1 was more prominent at loci lacking P300 (Fig-7A). Promoters displaying this motif included genes for *Stat6, Adamts1, Socs1*, and several interferon regulated genes (*Irf1, Irf9, Mx2, Il15, Ifit1*). These loci displayed STAT3 binding in samples from wt mice and STAT1 binding in datasets from *Il6*^−/-^ mice (Fig-7A). A limited number of genomic loci showed binding for both STAT1 and STAT3 (Fig-7A).

**Figure 7.**
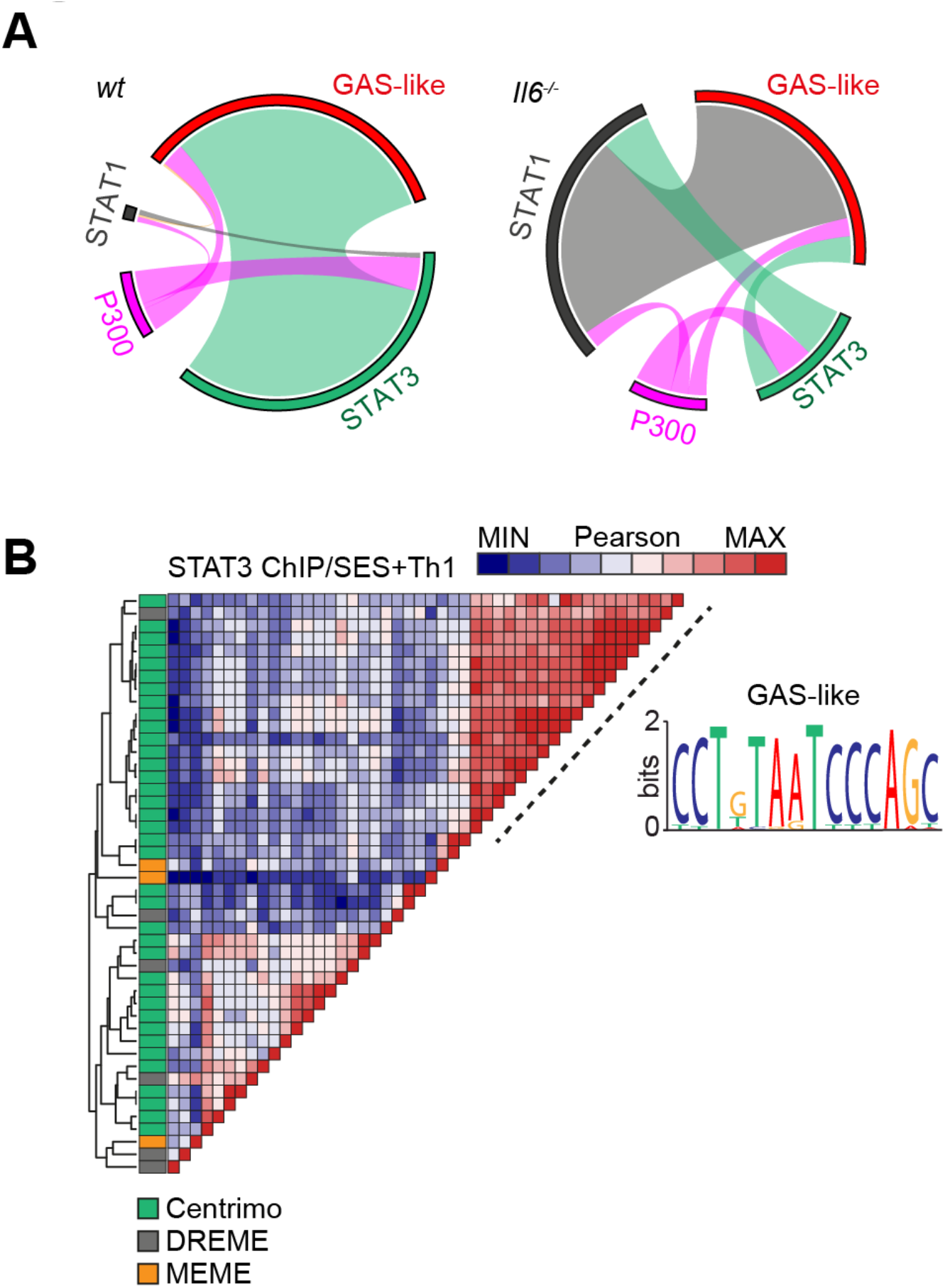
ChIP-seq provides evidence of STAT1-STAT3 cross-regulation. **(A)** Circos plots show the relative binding of STAT transcription factors to putative enhancers defined by either P300 binding (either *wt* or *Il6^−/−^* mice) or sites bearing homology to a *de novo* GAS-like motif identified by STAT1 and STAT3 ChIP-seq (Top). Binding to these sites was calculated using the Bedtools intersect algorithm. **(B)** Motif enrichment analysis of STAT3 ChIP-seq dataset from wt mice (SES+Th1). Pairwise comparison using the Pearson method was generated using Motif Alignment and Search Tool (MAST; MEME ChIP suite). Annotation shows the source algorithm of each motif (Centrimo (green), MEME (orange), and DREME (grey); MEME ChIP suite). A cluster is highlighted (hashed line) that maps to a sequence displaying homology with a GAS-like motif.

Combining *wt* and *Il6*^−/-^ mice datasets, motif analysis identified a centrally enriched sequence (CCTGTAATCCCAGC) with 90-95% identity to annotated GAS elements (MA0137.3, MA0144.2; Supplemental Figure-3B) in our STAT1 and STAT3 ChIP-seq data (Fig-7B). Given the conserved nature of this sequence, we used coordinate mapping to define genomic regions with proximity to the CCTGTAATCCCAGC motif in the murine genome. Here, sequence analysis showed the CCTGTAATCCCAGC motif to reside in short interspersed nuclear elements (SINE) classified by the RepeatMasker bioinformatic tool (Fig-8) (Ferrari *et al*, 2020). Analysis located STAT1 and STAT3 binding to a subset of SINE resembling B1 Alu elements (Fig-8A, Supplemental Figure-4). These elements display conserved architectures that include the CCTGTAATCCCAGC sequence (termed the GAS-Alu motif), residing close to a Pol-II A-box and flanked by consensus sites for T-bet and Runx3 (Fig-8B). Genomic DNA from SES-challenged mice showed no significant interaction of STAT transcription factors with the GAS-Alu motif. Thus, Th1 cells modify Jak-STAT cytokine signaling by re-directing STAT factors to genomic *Alu*-like retroelements.

**Figure 8.**
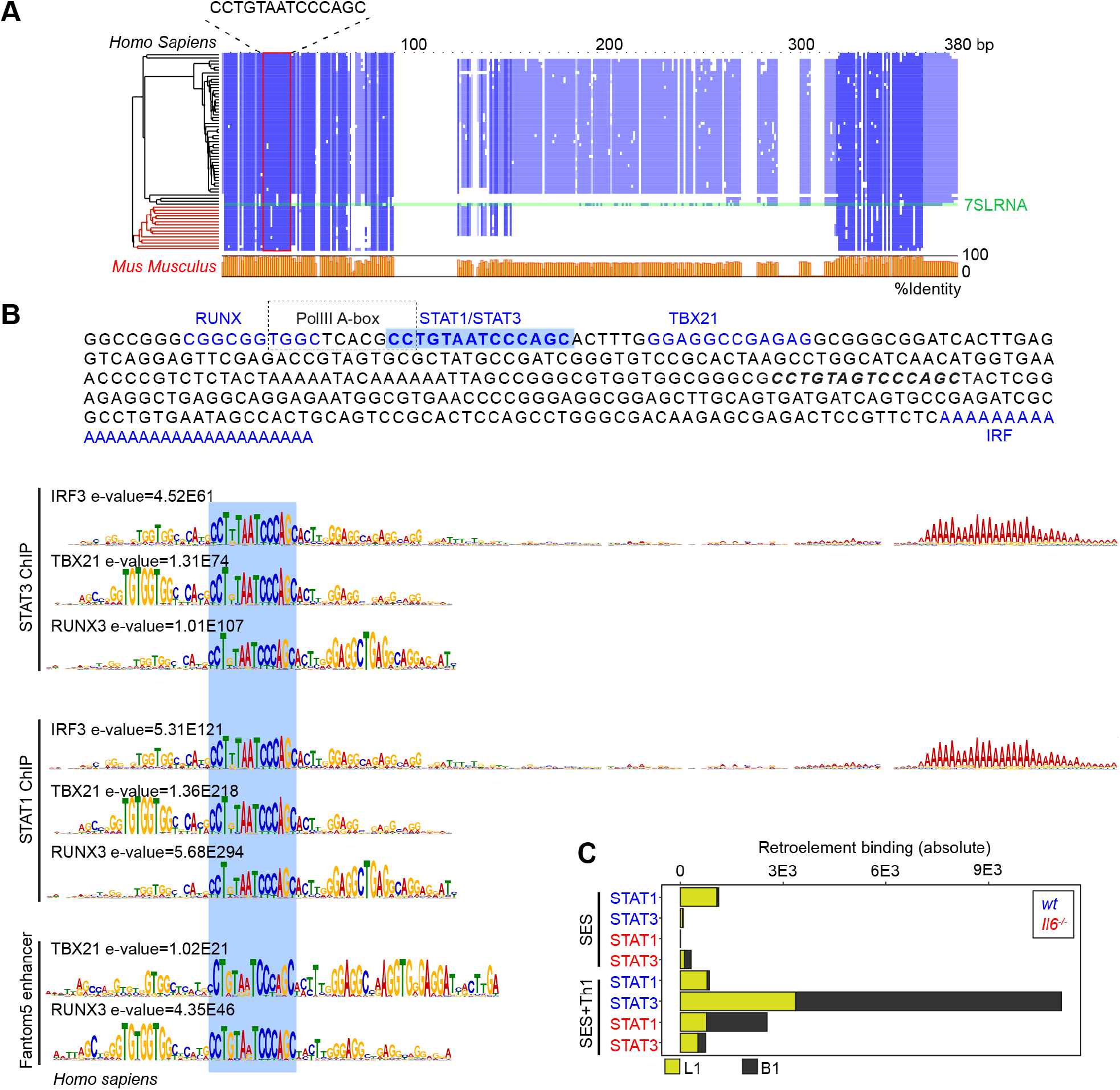
Identification of a GAS-like motif in Alu elements. **(A)** Multiple sequence alignment (Muscle; EBI) of human and mouse retroelement sequences downloaded from the dfam database (n=66 sequences). The Alu-GAS motif (CCTGTAATCCCAGC) identified by STAT1 and STAT3 ChIP-seq is located. Conserved regions are shown in blue, and a summary of the sequence identity is shown in orange (0–100%). **(B)** An annotated summary retroelement sequence is shown (top) locating representative secondary motifs (+/−150bp) relative to the Alu-GAS motif. Spaced motif analysis (Spamo; Meme ChIP suite) of these secondary sites are shown. **(C)** Quantitation of retroelement binding in ChIP-seq datasets based on repeatmasker annotations.

### The GAS-like motif identifies immune pathways linked to human physiology

Based on the sequence homology between murine and human Alu elements, we tested whether the GAS-Alu motif correlated with single nucleotide polymorphisms (SNP) in human disease (Fig-9A). From publicly available genome-wide association studies (GWAS), we identified GAS-Alu motifs in enhancer sequences designated by the FANTOM5 consortium. Our analysis revealed 8423 sequences, mapping to 2334 genes (Entrez) (Supplemental Figure-5A). Genes affiliated with the GAS-Alu motif were involved in various processes, including thrombopoietin (e.g., *PRKCB*) and VEGF (e.g., *PDGFC, ACTG2*) and integrin (e.g., *RAPGEF1*) signaling. Others regulate leukocyte signaling (e.g., *VAV1, CACNG3, PPP3CB*) and migration (ARHGAP8, ACTG2), and tissue turnover (e.g., *MYO10, ARHGEF19*). These included several mapped by our ChIP-seq of murine STAT1 and STAT3. For example, *ARHGEF19 (Arhgef19), COL5A (Col5a), MYO10 (Myo10), PPARG (Pparg), PRKCB (Prkcb), and RABGEF1 (Rabgef1)*.

**Figure 9.**
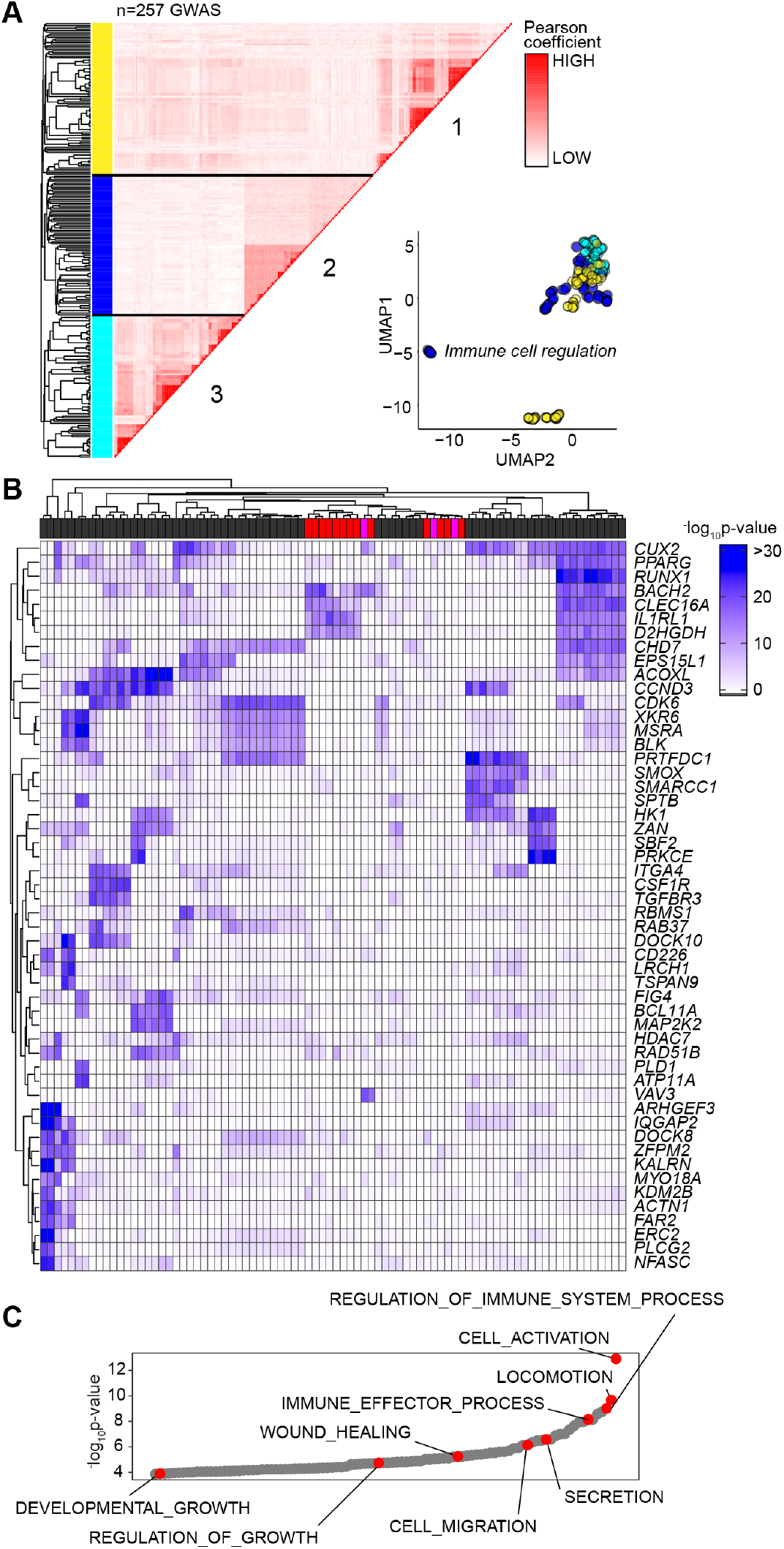
Association of GAS-Alu motif with human physiology. **(A)**Enrichment of GWAS signal in genes containing the GAS-Alu motif. MAGMA output files were compared against enhancers mapped by the FANTOM5 project (see ***Supplemental Figure 5***). For each GWAS, a vector was formed of the -log p-values of the motif-linked genes, and Pearson correlations calculated for each pair of p-value vectors. Analysis shows the 257 GWAS displaying significant enrichment for associations. UMAP distribution identifies GWAS datasets sharing common traits. Those with links to immune cell function (Cluster-2) are highlighted in Blue. **(B)**Heatmap of 52 gene targets displaying the top gene-wide p-values identified in Cluster-2 phenotypes. Horizontal bar colors designate GWAS phenotypes linked to immune cell regulation (grey; n=68); immunopathology (red; n=13); and other (pink; n=3). **(C)** Gene Ontology enrichment analysis of the genes identified in Panel-B (n=53 biological functions). Examples of processes are highlighted in Red.

Next, we downloaded GWAS summary statistics (n=2505) hosted by major repositories (NHGRI-EBI, CTGLab, NCBI) and tested for enrichment of GWAS signals in FANTOM5 genesets and gene signatures of IL-6 and IFNγ activity (available from msigdb; Broad Institute) using MAGMA (de Leeuw *et al*, 2015). Gene-wide significance of genes in these genesets were compared to a series of randomized and shuffled gene lists. Genes aligned to the HLA locus were excluded from our datasets to control against the high degree of linkage disequilibrium at these loci. Testing enrichment of GWAS signal in genes containing the GAS-Alu motif, MAGMA output files were compared against enhancers mapped by the FANTOM5 project. We identified 257 GWAS displaying enrichment of the GAS-Alu motif (Supplemental Figure-5B). These included GWAS linked with immune cell regulation, immune pathologies, including asthma, allergy and heart disease, and others linked with metabolism. A pairwise comparison (Pearson) of genes containing the GAS-Alu motif, based on their GWAS significance, is shown (Fig-9A). Here, hierarchical clustering of these GWAS datasets identified three clusters (Fig-9A). One of these clusters showed enrichment of IL-6 and IFNγ gene signatures and included genes involved in immune regulation and pathophysiology (Fig-9B & 9C). These include traits assigned as polygenic and monogenic disease variants (Vuckovic *et al*, 2020). Thus, the GAS-Alu motif classifies examples of Jak-STAT cytokine signaling in immune pathology.

## Discussion

Cytokines regulate transcriptional processes that maintain tissue homeostasis, protective immunity, and outcomes affecting inflammation-induced tissue injury (Borthwick *et al*, 2013; Duffield *et al*, 2013). In bacterial peritonitis, IL-6 and IFNγ are critical determinants of these outcomes (Catar et al., 2017; Fielding et al., 2014; Hurst et al., 2001; Jenkins et al., 2005; Jones et al., 2010; Krausgruber et al., 2020; McLoughlin et al., 2005; McLoughlin et al., 2003; Modur *et al*, 1997; Navarini et al., 2011; Robson et al., 2001; Schett et al., 2013; Xing *et al*, 1998; Yang *et al*, 2020). Here, IL-6 compromises tissue repair by supporting the expansion of pro-fibrotic IFNγ-secreting CD4^+^ T-cells (Fielding et al., 2014). While these cytokines rely on STAT1 and STAT3 signaling (Fielding et al., 2014; Fielding et al., 2008; Jenkins et al., 2005; McLoughlin et al., 2005), it is unclear how these transcription factors coordinate anti-microbial host immunity, tissue scarring, and fibrosis. To understand their involvement in these processes, we tracked the stromal activity of STAT1 and STAT3 transcription factors in peritoneal tissues following an inflammatory challenge. Our analysis shows that IFNγ-secreting CD4^+^ T-cells alter the transcriptional output of IL-6 by channeling STAT transcription factors to previously unoccupied genomic loci.

Jak-STAT signaling is complex and includes an interplay between individual STAT transcription factors (Costa-Pereira et al., 2002; Ernst *et al*, 2008; Hirahara et al., 2015; Jones et al., 2013; Olbrich & Freeman, 2018; Qing & Stark, 2004; Villarino et al., 2017; Yang et al., 2020; Yang *et al*, 2011). For example, patients with STAT1 gain-of-function or STAT3 loss-of-function mutations often display similar clinical features. These include increased susceptibility to infections at barrier surfaces, eczema-type rashes, and bowel perforations (Abusleme *et al*, 2018; Goel *et al*, 2018; Liu *et al*, 2011; Luis & Calva-Mercado, 2018; Milner *et al*, 2015; Olbrich & Freeman, 2018; Toubiana *et al*, 2016). These clinical phenotypes are also present in patients lacking IL6R or mice lacking *Il6* and often reflect the role of STAT3 in innate immunity (Holland *et al*, 2007; Kano *et al*, 2003; Kopf *et al*, 1994; Matsukawa *et al*, 2003; Ramsay *et al*, 1994; Spencer *et al*, 2019; Welte *et al*, 2003). Consistent with these findings, our analysis shows that IL-6 controls stromal activities that enhance the phagocytic properties of infiltrating neutrophils. The inclusion of Th1 cells enhances this anti-microbial response, contributing to an increase in various interferon-responsive genes supporting host immunity. During acute resolving inflammation, a limited number of IFNγ-secreting CD4^+^ T-cells are present in the peritoneal cavity (Fielding et al., 2014; Jones et al., 2010). This number significantly increases as a response to repeated peritonitis, and their retention leads to compromised tissue homeostasis and fibrosis following activation (Fielding et al., 2014; Jones et al., 2010). Our analysis reveals that this change in inflammatory status impacts various cellular processes that include an alteration in IL-6 bioactivity. These data point towards a transcriptional interplay between IL-6 and IFNγ that impacts how STAT transcription factors shape inflammatory outcomes.

Studies of cancer cells and ex vivo stimulated T-cells detail how cross-regulation between STAT1 and STAT3 modulates target gene expression (Costa-Pereira et al., 2002; Hirahara et al., 2015; Hong et al., 2002; Jones et al., 2013; Twohig, 2019). Our studies suggest that IFNγ secretion by Th1 cells may facilitate such interactions in stromal tissues following inflammatory activation. In this context, Th1 cells directly impacted the transcriptional output of IL-6 and were responsible for directing STAT1 and STAT3 to previously latent enhancers. Genes affiliated with these loci contribute to tissue remodelling, fibrosis, solute transport, membrane permeability, and hypoxia. Thus, our data points towards an agonist-specific repertoire of latent enhancers employed to sense and interpret changes in the tissue microenvironment (Ostuni *et al*, 2013). For example, transcription factors linked with myeloid cell development (e.g., PU.1) often instruct the binding of NF-κB, AP-1, and interferon response factors to the genome (Natoli *et al*, 2011). Our results are, therefore, consistent with theories that genes linked by roles in related biological functions commonly share similar mechanisms of transcriptional control (Tong *et al*, 2016). In this regard, a close inspection of the DNA sequences enriched for STAT transcription factor binding identified a conserved motif in *Alu*-like retroelements (Chen & Yang, 2017; Vassetzky & Kramerov, 2013). Retroelements are endogenous components of eukaryotic genomes. They support non-allelic recombination, polyadenylation, alternative splicing, and the transcription of gene-rich regions (Chen & Yang, 2017; Tajnik *et al*, 2015). Significantly, retroelements possess consensus binding motifs for various transcription factors and often display evidence of DNA methylation, suggesting an involvement in gene regulation (Polak & Domany, 2006; Xie *et al*, 2009). Functional genomic studies in cell lines of stromal or immune cell origin demonstrate the binding of basic leucine zipper transcription factors, the aryl hydrocarbon receptor and other transcriptional regulators to retroelements (Lu *et al*, 2020; Roman *et al*, 2008; Sun *et al*, 2018). Our analysis showed that IL-6 signaling, in association with IFNγ-secreting Th1 cells, promotes STAT3 binding to Alu sequences. STAT3 binding was, however, lost in *Il6* deficiency. Instead, these same sites showed STAT1 binding. Thus, Alu-like retroelements may represent sentinels of transcriptional cross-regulation in stromal tissues.

What is the significance of STAT transcription factor binding to *Alu* sequences? Are these interactions relevant to human disease or treatment responses to tocilizumab, tofacitinib, and others? Addressing these questions is challenging due to the complexities of mapping repetitive DNA elements. Our data imply a link between *Alu*-like sequences and tissue pathology. Moreover, GWAS commonly identify human *Alu* polymorphisms linked with interferonopathies or diseases characterized by alterations in STAT1 activity (Hung *et al*, 2015; Payer *et al*, 2017; Volkman & Stetson, 2014; Zeng *et al*, 2014). Our analysis of human GWAS datasets revealed several gene targets also identified in mice treated with SES and Th1 cells. These data support a role for epigenetic modifiers that regulate the accessibility of transcription factors to specific enhancers under certain inflammatory settings or disease processes. Future evaluation of these events will open new opportunities to understand how cytokine cues are interpreted or fine-tuned to direct physiology or pathophysiology.

## Acknowledgements

This manuscript is dedicated to Dr Javier Uceda Fernandez, a dearly loved friend and colleague who was tragically taken from us on the 29th of August 2018. Kidney Research UK (Reference RP-024-20160304 awarded to SAJ, DF, NT, GWJ), Versus Arthritis (Reference 20770, 19796, 20305 awarded to SAJ, VOD, NMW & GWJ), and the National Health and Medical Research Council of Australia (to BJJ) provided grant support for this project. JUF was recipient of a la Caixa PhD Studentship administered through the British Council, and BCC is supported by a PhD studentship from the Systems Immunity University Research Institute at Cardiff. JL is recipient of a Rutherford Fellowship Grant. PRT is recipient of a Wellcome Trust Investigator Award (Reference 107964/Z/15/Z) and receives funding through the UK Dementia Research Institute. Bioinformatic analysis was developed with support from the Systems Immunity University Research Institute in Cardiff.

## Materials and methods

### Animals

All procedures were performed under UK Home Office project license P05D6A456. Inbred wild type C57BL/6 male mice were purchased from Charles River UK. IL-6-deficient (*Il6*^−/-^) (Kopf et al., 1994) were bred under approved UK Home Office guidelines in Cardiff University. The *gp130*^Y757F:Y757F^ and *gp130*^Y757F:Y757F^:*Stat3*^+/−^ mice have been previously described (Jenkins et al., 2005). Experiments were approved by the Animal Ethics Committee and included genetically matched *gp130*^+/+^ littermate controls. All experiments were performed using age-matched 8-12 weeks old mice.

### SES-Induced Peritoneal Inflammation

A lyophilized cell-free supernatant prepared from *Staphylococcus epidermidis* (SES), whose activity had been standardized using an in vitro cell-based CXCL8 bioassay, was used to induce acute peritoneal inflammation (Hurst et al., 2001). Mice were administered (i.p.) with a defined dose of SES (500μl). Soluble gp130Fc was added (i.p.) to wt mice as indicated. At 3 and 6 hours post-inflammatory challenge, mice were sacrificed and the peritoneal cavity lavaged with 2ml ice cold PBS. The peritoneal membrane was harvested at the same time points.

### Transfer of Th1 Cells

To replicate events promoting peritoneal fibrosis some mice receiving SES were also simultaneously administered with either naïve CD4^+^ T-cells or CD4^+^ T-cells conditioned ex vivo towards a Th1 phenotype. Briefly, splenic naïve CD4^+^ T-cells (CD4^+^CD25^−^CD44^lo^CD62L^hi^) were flow sorted and place in coated plates with anti-CD3e (145-2C11) and 5μg/ml soluble anti-CD28 (37.51) antibodies. Cells were cultured for 4 days in the presence of 20ng/ml murine recombinant IL-12 (R&D systems; 419-ML). The proportion of IFNγ-secreting CD4^+^ T-cells (Th1 cells) was determined by intracellular flow cytometry using antibodies against to CD4 (RM4-5), IFNγ (XMG1.2), IL-17 (TC11-18H10.1), IL-4 (11B11) and IL-13 (eBio13A). Based on this analysis, T-cells were washed in ice cold PBS, re-suspended in a 500μl aliquot of PBS reconstituted SES, and administered via the intraperitoneal route. Here, a cell concentration of 5-10 ×10^5^ CD4^+^ T-cells was administered to reflect the proportion of Th1 cells recruited to the peritoneal cavity under acute SES challenge. Where indicated, control mice were administered the same number of sorted naïve (Th0) CD4^+^ T-cells. Changes in the inflammatory infiltrate were analyzed by direct counting (Coulter Z2, Beckman Coulter), differential cell counting and flow cytometry approaches using antibodies against defined leukocyte subsets. The peritoneal membrane was harvested at 3 and 6h after the injections.

### Fluorescent labelling of bacteria

An inoculum of *Staphylococcus epidermidis* ATCC 12228 (5×10^8^ cfu/mouse) was prepared from log phase cultures under sterile conditions. The suspension was centrifuged, and bacteria labelled for 20 min at 37°C in pre-warmed PBS containing Cell Trace Far Red (CT-FR) (Life Technologies) (1 μM or 8 μM for *ex vivo* and *in vivo* experiments, respectively). For ex vivo experiments, bacteria were serum-opsonized while, for in vivo experiments were centrifuged and washed 3 times in PBS, before resuspending in sterile PBS.

### Ex vivo neutrophil effector function assay

Whole blood was collected by cardiac puncture into tubes containing 5mM EDTA. Samples were diluted 1:10 and washed 3 times in ice cold PBS. Cells were resuspended in serum-free RPMI 1640 containing 5 μM of 3’-(p-aminophenyl) fluorescein (APF) and incubated for 30 min at 37°C. APF-loaded cells were split into 100 μl aliquots and cultured at 37°C with an equal volume of pre-warmed, opsonized Cell Trace Far Red (CT-FR; 1 μM) labelled *S. epidermidis*. Cells were incubated for set time intervals (0-30 min) and transferred to an iced water bath prior to preparation for flow cytometry using a Beckman Coulter Cyan-ADP flow cytometer.

### In vivo neutrophil effector function assay

Mice were administered (i.p.) with Cell Trace Far Red (CT-FR; 8 μM) labelled S. epidermidis. An independent group of *Il6*^−/-^ mice received a dose of CT-FR labelled *S. epidermidis* together with the IL-6-sIL-6R chimeric fusion protein HDS (50 ng/ml). Six hours after bacterial challenge, the peritoneal cavity was lavaged with 2 ml of RPMI 1640 containing 5 μM of 3’-(p-aminophenyl) fluorescein (APF). Neutrophil phagocytosis and respiratory burst activity were examined by flow cytometry using a Beckman Coulter Cyan-ADP flow cytometer and analyzed using *Summit* (software v4.3, Beckman-Coulter) or FlowJo 10 (TreeStar).

### Imaging flow cytometry analysis

Lavaged neutrophils were immune-stained for Ly6G (1A8), cells were resuspended in 100 μl of sterile PBS, and events (>8000 events per sample) were acquired at a low imaging rate and 60X amplification using the Amnis ImageStream^®^X Mark II Imaging Flow Cytometer (Amnis). Neutrophils were gated according to Ly6G staining. Phagocytic activity was expressed as a Phagocytic Index, which reflects the number of bacteria ingested by an individual Ly6G^+^ neutrophil during the incubation period. To evaluate the phagocytic index of neutrophils undergoing phagocytosis, the Application “Spot Counting” in ImageStream software IDEAS (Amnis) was used. Spots were counted based on the distribution of CT-FR bacterial staining. The efficiency of phagocytic uptake was quantified by examining cells displaying either a low (1-2 counts) or a high number (3 counts or more) of ingested bacteria.

### Immunoblotting of peritoneal tissues

Protein was extracted from frozen peritoneal biopsies using ice-cold lysis buffer. Samples were pre-cleared of cellular debris before separation by SDS-PAGE and immunoblotting with specific antibodies against STAT1, STAT3 and tyrosine phosphorylated forms of STAT1 (pY-STAT1) and STAT3 (pY-STAT3) (Jenkins et al., 2005). Immuno-labelled proteins were detected using either the enhanced chemiluminescence (ECL) detection system (Amersham Biosciences) or Odyssey Infrared Imaging System (LI-COR, Lincoln, New England) with the appropriate secondary antibodies as per the manufacturer’s instructions.

### RNA-seq

Peritoneal membrane sections (80mg tissue) were dissociated in 1ml buffer RLT (QIAGEN) supplemented with β-mercaptoethanol (1:100 v:v) using a handheld electric homogeniser (Benchmark Scientific). Lysate was diluted 1:3 in distilled water and digested in 0.2mg/ml proteinase-K (Invitrogen; 25530049) for 10 minutes at 55°C. Lysate was cleared and RNA precipitated in 70% ethanol. Total RNA was extracted using the RNeasy Mini kit (QIAGEN) following the manufacturer’s instructions. 2-4mg of mRNA was processed to generate the libraries. Cytoplasmic, mitochondrial, and ribosomal RNA was depleted using the Ribominus transcriptome isolation kit (Ambion; K155001). Libraries were prepared using the RNA-seq kit v2 (Life technologies; 4475936) and sequencing on an ion torrent (Thermo Fisher).

### Chromatin Immunoprecipitation (ChIP)-seq

Excised peritoneal membranes were immediately frozen in liquid nitrogen and stored at −80°C until use. Tissues were diced and ground to a fine powder with intermittent addition of liquid nitrogen. Genomic DNA was extracted, crosslinked, and fragmented by sonication prior to treatment with 2μg/ml of anti-STAT1 (sc-592, Santa Cruz Biotechnology), anti-STAT3 (sc-482, Santa Cruz Biotechnology), anti-P300 (05-257, Millipore), or isotype control antibodies. Immunoprecipitation was conducted overnight at 4°C under continuous gentle agitation. Antigen-antibody complexes were captured using Protein-A/G magnetic beads, washed and DNA fragments liberated following treatment with proteinase-K and extraction with phenol-chloroform. Biological repeats from 3 independent tissue extracts were pooled and concentrated before the library preparation and next generation sequencing. ChIP libraries were prepared according to the manufacturer’s instructions (Illumina Truseq DNA ChIP kit; RS-122-2001). Size selection (200-400bp) was determined using a Blue Pippin (Sage Science) system employing 2% agarose cartridges (Sage Science; BDF2003). Appropriate library size distribution was confirmed using an Agilent 2100 bioanalyser and final concentrations determined using Qubit (Invitrogen). Libraries were sequenced on an Illumina HiSeq4000.

### ATAC-seq

Excised peritoneal membranes were immediately frozen in liquid nitrogen and stored at −80°C until use. Tissues were diced and ground to a fine powder with intermittent addition of liquid nitrogen. Omni-ATAC seq was performed as described previously (Corces *et al*, 2017). Briefly, 100,000 nuclei per sample were isolated using an iodixanol gradient and ATAC-seq was subsequently performed according to the original protocol (Buenrostro *et al*, 2013) using Nextera DNA Sample preparation kit (Illumina, FC-121-1030). After library amplification, DNA was isolated using Qiagen MiniElute kit and size selection was performed using Blue Pippin (Sage Science) system employing 2% agarose cartridges (Sage Science; BDF2003). Libraries were sequenced on an Illumina HiSeq4000.

### Quantification and Statistical Analysis

No randomization and no blinding were used for the animal experiments. Whenever possible, the investigator was partially blinded for assessing the outcome. All data were analyzed using Prism 8 (GraphPad Prism, La Jolla, CA). Information on the types of Statistical methods used, the sample size and number of repetitions are listed in the Legends.

### RNA-seq data analysis

Raw fastq were mapped using Torrent SuiteTM to the *mm10* reference genome and counts were assigned to transcripts using featureCounts (Liao *et al*, 2014) with the GRCm38.84 Ensembl gene build GTF. Differential gene expression analyses used the DESeq2 package (Love *et al*, 2014). Genes were discarded from the analysis differential expression failed to be significant (significance: adj.pval < 0.05, Benjamini-Hochberg correction for multiple testing). Differentially regulated genes were uploaded into Ingenuity Pathway Analysis (IPA) (QIAGEN) for functional analysis.

### ChIP-seq data analysis

Between 40-70M reads were obtained for each sample. These were trimmed with Trim Galore (https://www.bioinformatics.babraham.ac.uk/projects/trim_galore/) and assessed for quality using FastQC (https://www.bioinformatics.babraham.ac.uk/projects/fastqc/). Reads were mapped to the mouse mm10 reference genome using bwa (Li & Durbin, 2009). Peaks were called using macs2 (Zhang *et al*, 2008), using the BAMPE model (adj.pval 0.05).

### ATAC-seq analysis

**P**aired-end reads were processed with Trim Galore (https://www.bioinformatics.babraham.ac.uk/projects/trim_galore/) and assessed for quality using FastQC (https://www.bioinformatics.babraham.ac.uk/projects/fastqc/) prior to mapping to the mouse mm10 reference genome (Li & Durbin, 2009). Peaks were called using macs2 and the BAMPE model (adj.pval 0.05) (Zhang et al., 2008). Differential open-region analysis used Diffbind in Bioconductor (http://bioconductor.org/packages/release/bioc/vignettes/DiffBind/inst/doc/DiffBind.pdf). Histograms depicting distance from TSS were generated using the ChIPseeker R-package and show the peak count fraction.

### FANTOM5 enhancer analysis

Sequences annotated as enhancers by the FANTOM5 consortium were downloaded from Slidebase (250bp pad) (http://slidebase.binf.ku.dk). Motif occurrences (n=8423, p<0.00001) in these sequences were identified using the FIMO algorithm (Meme ChIP-suite; http://meme-suite.org/tools/fimo). Genes in proximity to the identified sequences (defined as 2kb upstream or downstream) were mapped to the ensembl GRCh38 (hg38) build and visualized using the ClueGO cytoscape pluggin). Genome-wide motif occurrences were visualized with RIdeogram R-package (https://cran.r-project.org/web/packages/RIdeogram/vignettes/RIdeogram.html).

### Motif identification

Sequences under ChIP peaks (q<0.05) were obtained using the bedtools getfasta command against the ensemble *mm10* genome build. Sequence fasta files were uploaded to Meme ChIP (http://meme-suite.org/tools/meme-chip) and searched against the murine HOCOMOCO (v11 CORE) and Eukaryote DNA databases. The de novo GAS-like motif was enriched by both Centrimo and Meme algorithms.

### Spaced motif analysis

Analysis was conducted with Spamo (Meme-ChIP-suite) using default parameters. Secondary motifs occur within 150bp of the user-provided primary motif (GAS-like). All secondary motifs are referenced in the HOCOMOCO (v11 CORE) database. Input sequences, including a 250bp pad, were derived from the ChIP-seq data or enhancer annotations in FANTOM5.

### Multiple Sequence Alignment

Consensus sequences (Hidden Markov Models) corresponding to murine and human Alu-family transposable elements were downloaded from the Dfam database (https://dfam.org/) - Mouse; B1_mm, B1_mus1, PB1D11, B1_Mur2, B1_Mur4, PB1, B1_Mur1, B1_Mur3, PB1D10, B1_Mus2, B1_F1, PB1D7, PB1D9, B1F2, B1F, B2_Mm1a, B2_Mm1t, B2_Mm2, B3, B3A, B4, B4A, ID_B1, Human; AluY, AluSc, AluJB, AluJo, AluJr, AluJr4, AluSc5, AluSc8, AluSg, AluSg4, AluSg7, AluSp, AluSq, AluSq10, AluSq2, AluSq4, AluSx, AluSx1, AluSx3, AluSx4, AluSz, AluSz6, AluYa5, AluYa8, AluYb8, AluYb9, AluYc, AluYc3, AluYd8, AluYh9, AluYk11, AluYk12, AluYk4, AluYg6, AluYk3, AluYm1, AluYk2, AluYe6, AluYi6, AluYe5, AluYi6_4d, AluYf1, AluYh3, AluYj4, AluYh7, FAM, FLAMA, FLAMC, FRAM. Sequences were aligned using MUSCLE (European Bioinformatics Institute) using the default ClustalW output and visualized in Jalview.

### Gene Set Enrichment Analysis (GSEA)

A ranked gene list was prepared for each dataset using the differential gene expression analysis. Using the GSEA pre-ranked function, enrichment profiles were generated against the biological processes (C-5; 7573 gene sets) gene ontology database. For visualization, GSEA output files were loaded into Cytoscape using the enrichment-map plugin. For small gene sets, ontology enrichments were performed using either the mSigdb overlap tool (broad institute) or the Metascape online tool. Network aesthetics (e.g., color, spacing) were modified in adobe illustrator. Network statistical thresholds are below p<0.01, q<0.01.

### Repeatmasker overlaps

Alu and L1 sequence coordinates were derived from the repeatmasker definitions downloaded from the UCSC table browser (mm10). Repeats overlapping STAT binding were identified using the bedtools intersect algorithm. The absolute number of Alu and L1 sequences overlapping each ChIP dataset was calculated in R studio and plotted using the circlize package.

### Visualization and annotation

Heatmap visualization in Morpheus (broad institute) and Pheatmap (R package) using colours defined in the viridis and R Color Brewer packages. Figures were prepared using the ggplot2 R package (R package), graphpad prism8, and arranged in Adobe Illustrator. Data filtering for complex heatmaps was achieved using Dplyr (R package). IL-6 regulated genes were defined as those meeting the statistical threshold (*Il6*^−/-^ vs wt padj<0.05 with a log2 fold change greater than 1.75 or less than −1.75) in at least one experimental condition. STAT regulated genes were similarly described as those with ChIP signal under the statistical threshold (q<0.05) in at least one experimental condition.

### Generation of gene sets for MAGMA analysis

Sequences annotated as enhancers by the FANTOM5 consortium were downloaded from Slidebase (http://slidebase.binf.ku.dk/) including a 200bp pad. To identify motif occurrences, these sequences were entered to the FIMO web server (Meme ChIP suite) together with the MEME-formatted motif extracted from our ChIP-sequencing analyses. Identified gene sets were derived by mapping coordinates to the HG19 reference genome (genes within 2kb) using PAVIS (https://manticore.niehs.nih.gov/pavis2/). In parallel, FANTOM5 sequences containing Alu sequences were defined by intersecting (bedtools) FANTOM5 coordinates with the Repeatmasker database (UCSC table browser). Coordinates were mapped to genes using PAVIS. Randomised control gene sets were derived as follows. (1) FIMO output coordinates were ‘shuffled’ across FANTOM sequences or genome wide (Hg19 reference genome) using the Bedtools shuffle feature. (2) Random bed files (8000x 500bp sequences) were generated using the Bedtools Random against either the FANTOM5 enhancer sequences or genome wide (hg19). Three gene sets were generated for each control. Hallmark gene sets were downloaded from msigdb (broad Institute). Genes mapping to the MHC locus were downloaded from the UCSC table browser. This gene list was used to filter MHC-locus genes from all gene sets prior to MAGMA analysis (Dplyr R-package).

### MAGMA

GWAS summary statistics (Hg19) were downloaded using FTP links supplied by the GWAS atlas (https://atlas.ctglab.nl/). After re-formatting for compatibility with MAGMA, summary statistics were mapped to genes using the build-37 gene locations file (NCBI37.3.gene.loc). Gene results files were next generated for each summary statistic. The GWAS atlas was used to refence N (number of study participants) for each study. Finally, gene set analysis was performed against gene sets generated as described above. Results files were read into R and relevant columns extracted using R base functions. For genes, p-values were extracted from MAGMA output files ending genes.sets.out. For gene sets, p-values were extracted from output files ending.gsa.out. P-values were merged into a single dataframe for correlation analysis (R-base function) or filtering (Dplyr R-package). Heatmaps were generated using the Pretty heatmap R package (Pheatmap).

### Data and Code Availability

RNA-seq, ChIP-seq and ATAC-seq datasets reported in this paper have been deposited in ArrayExpress under accession code EMBL-EBIE-MTAB-10087. The authors declare that all relevant data supporting the findings of this study are available on request. R scripts for performing the main steps of analysis are available from the Lead contact on reasonable request.

