## Supplemental data for "Th1 cells alter the inflammatory signature of IL-6 by channeling STAT transcription factors to *Alu*-like retroelements"

### Supplementary Figures & Legends

#### Supplemental Figure-1

**Gene Set Enrichment Analysis of RNA-seq data from SES challenged mice (A)** Enrichment map (Gary Bader; University of Toronto) visualization of Gene Set Enrichment Analysis (GSEA). GSEA was performed independently for each dataset using a ranked gene list ($Log_2FC$) against the molecular signatures database (msigdb) biological processes (C5) reference set.

#### Supplemental Figure 2

**Regulation of gene expression by IL-6 (A)** ELISA measurement of cytokines in lavage fluid following SES challenge. Data (mean ± SEM; n=4) is shown for mice treated with SES alone or in combination with Th1 cells. **(B)** Global transcriptomic analysis as in Figure-1, incorporating data from *Il6*$^{-/-}$ mice. The dataset is expanded to 1194 genes passing statistical thresholds in at least one experimental condition. **(C)** Pairwise comparison of transcriptomic data presented in Panel-B. **(D)** Volcano plots showing differential expression analysis (LIMMA) of transcriptomic data from wt and *Il6*$^{-/-}$ mice (*Il6*$^{-/-}$ vs *wt*). **(E)** Unsupervised clustering of omental stromal cells using publicly available scRNA-seq (GEO: GSM4053741; see Jackson-Jones *et al*, 2020). (Top Panels) The UMAP visualization shows cells classified as fibroblasts (blue) and mesothelial cells (red). The Top-10 genes identifying the classification of these cells is shown as a heatmap. (Bottom Panels) The heatmap shows a series of IL-6 regulated gene signatures (left) and their corresponding expression in the GEO dataset. UMAP visualizations are shown to two examples (*Angptl4*, *Serpina3n*). **(F)** Flow cytometric analysis of effector function of circulating neutrophils isolated from the peripheral blood of *wt* and *Il6*$^{-/-}$ mice. Measurements were recorded over a 30-minute time course (mean ± SEM; n=3). **(G)** Flow cytometric analysis of infiltrating neutrophils. Mice were challenged (i.p.) for 6 hours with DDAO-labelled *Staphylococcus epidermidis* ($5x10^8$ cfu). Cells were recovered by lavage, loaded with APF, and gated for Ly-6B$^{hi}$Ly-6G$^{hi}$ cells. The proportion of cells displaying neutrophil effector activities is shown (mean ± SEM; n=4). **(H)** Imaging flow cytometry of infiltrating neutrophils recovered from mice challenged with DDAO-labelled *Staphylococcus epidermidis* ($5x10^8$ cfu). Representative images are shown for cells recovered from wt, and *Il6*$^{-/-}$ mice and *Il6*$^{-/-}$ mice treated with an IL-6-sIL-6R fusion protein (50ng/mouse).

#### Supplemental Figure 3

**Analysis of STAT transcription factor activation following SES challenge (A)** Immunoblot analysis of STAT1 and STAT3 activity in peritoneal tissue extracts from SES challenged *wt*, *gp130*$^{Y757F:Y757F}$, and *gp130*$^{Y75F7:Y757F}$*:Stat3*$^{+/-}$ mice. Temporal changes in tyrosine phosphorylated STAT1 and STAT3 (pY-STAT1, pY-STAT3) are compared against total STAT1 and STAT3. Note, immunoblotting shown for STAT3 in *wt* and *gp130*$^{Y757F:Y757F}$ mice was previously presented in McLoughlin et al (2005) and is used as a control in this analysis. **(B)** GAS-like motif enriched in STAT1 and STAT3 ChIP-seq datasets (SES+Th1 condition). Shown are alignments (TomTom; Meme ChIP suite) to canonical GAS motifs (MA137.1; MA0137.2; MA137.3; MA144.1; MA0144.2). Significant alignment ($p<0.05$) can be achieved by breaking down the sequence into constituent parts (top right). **(C)** Histograms showing the number of peaks mapping to genomic features (ChIP-seq).

**Supplemental Figure 4**

**Enrichment of B1-like sequences in ChIP-seq datasets.** Motif sets generated by Meme analysis of ChIP-seq data were aligned to canonical B1 and Alu sequences downloaded from the Dfam Catalogue (n=66) using the Motif Alignment and Search Tool (MAST; Meme ChIP suite).

**Supplemental Figure 5**

**Bioinformatic pipeline underpinning the analysis of GWAS datasets. (A)** Flow chart summarizing the workflow; (1) instances of the GAS-Alu motif were identified across Fantom5 enhancers using FIMO (Meme ChIP suite); (2) Genes linked to these sites were mapped against the Hg19 reference genome; (3) Genes corresponding to the MHC locus were removed prior to MAGMA gene set enrichment analysis against 2506 GWAS summary statistics downloaded from major repositories (EBI, CTGLAB, NCBI); (4) P-values were extracted from MAGMA output files and correlated (Pearson). **(B)** Top 20 enriched phenotypes for the GAS-Alu motif linked gene set. Gene sets were filtered in R so that the highest scoring GAS-Alu motif-linked phenotypes are shown. Heatmap is clustered using the Euclidean method. The x-axis shows gene sets used for the analysis; Hallmark gene sets (msigdb); shuffled and randomized controls; FIMO mapped GAS-Alu motifs in Fantom5 enhancer sequences (motifs); Repeatmasker elements mapping to Fantom5 sequences (Fantom5_alu). HLA-mapped genes were removed prior from all gene sets to MAGMA enrichment analysis.

### Supplemental Figure-1

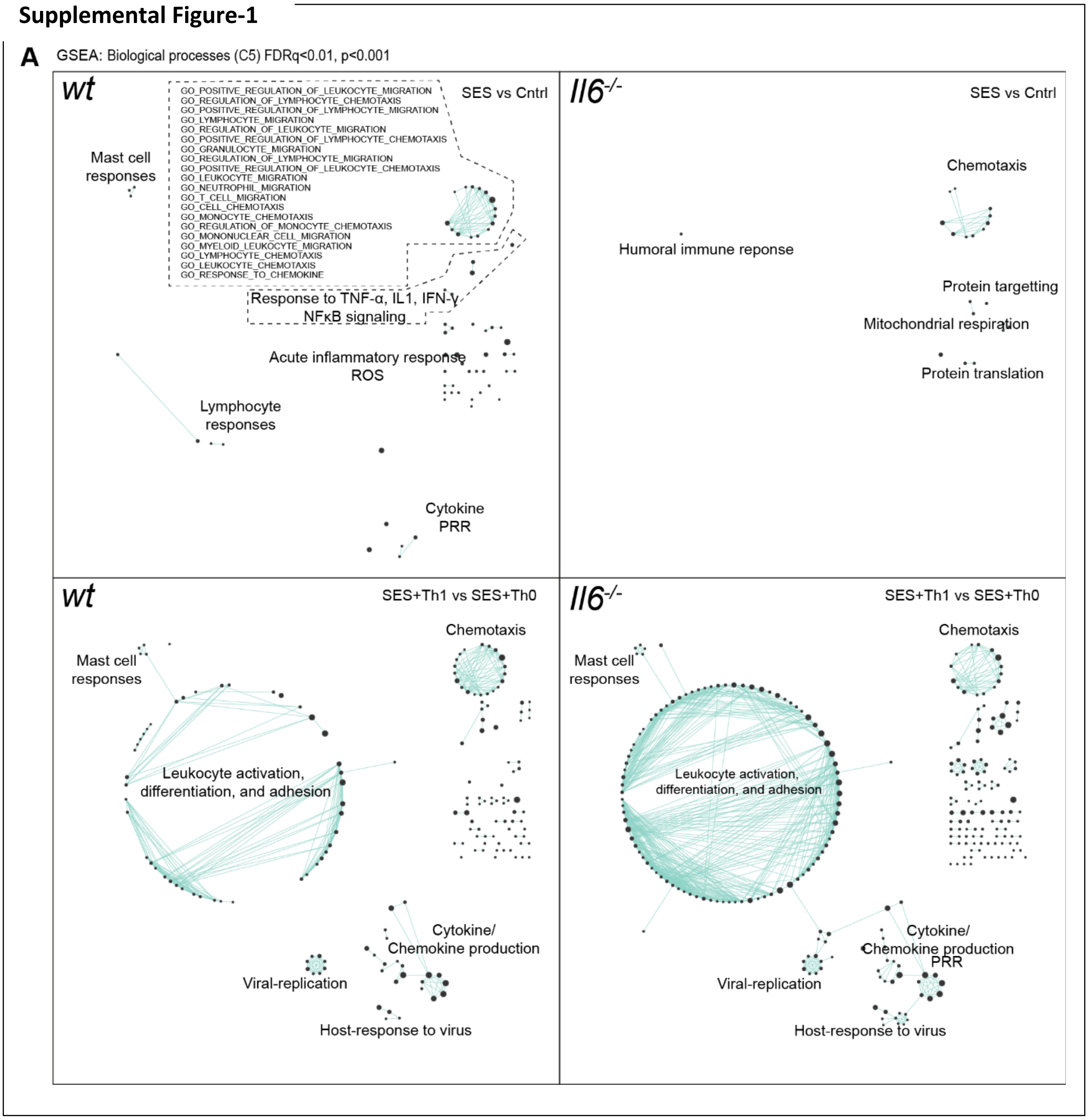

### Supplemental Figure-2

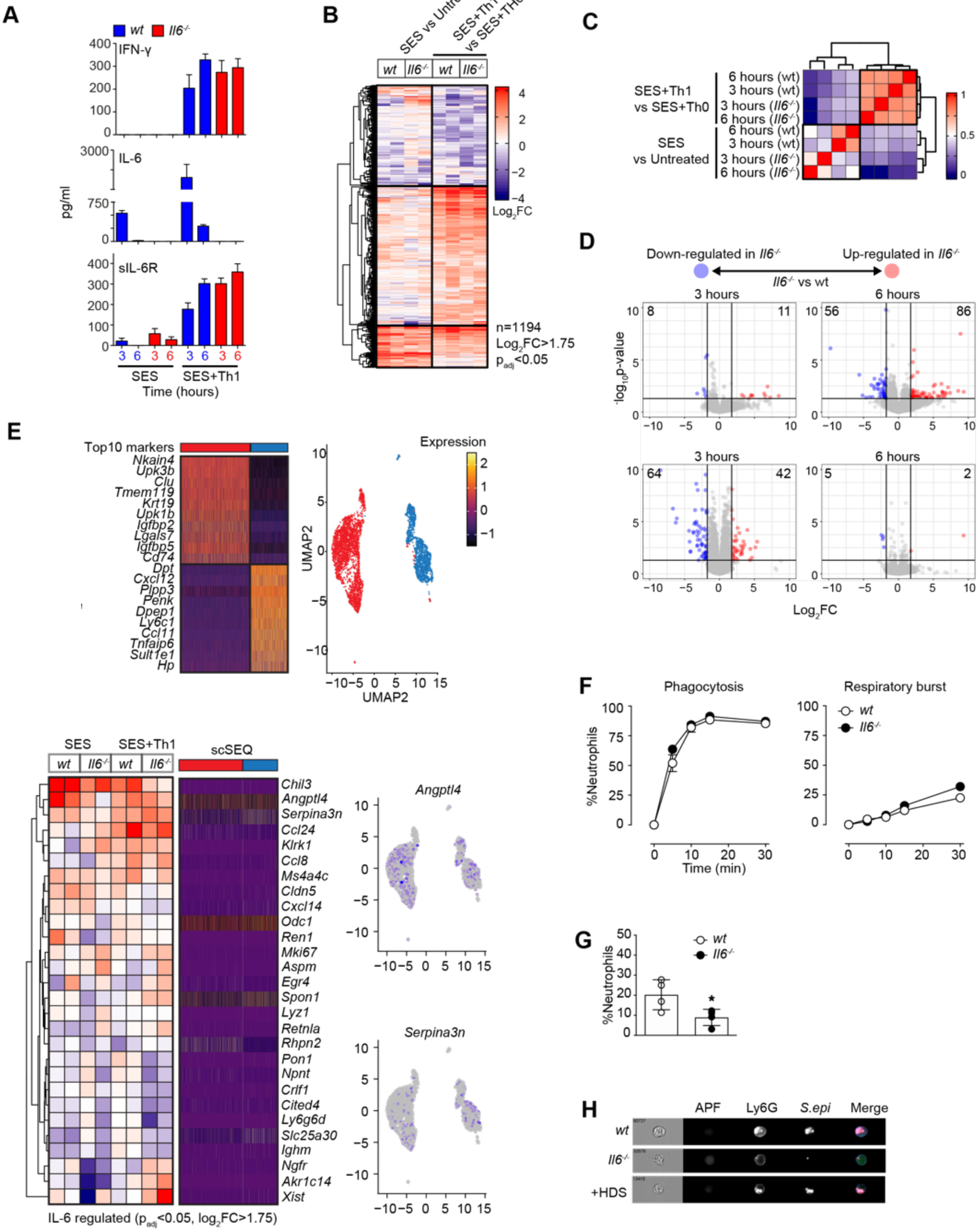

### Supplemental Figure-3

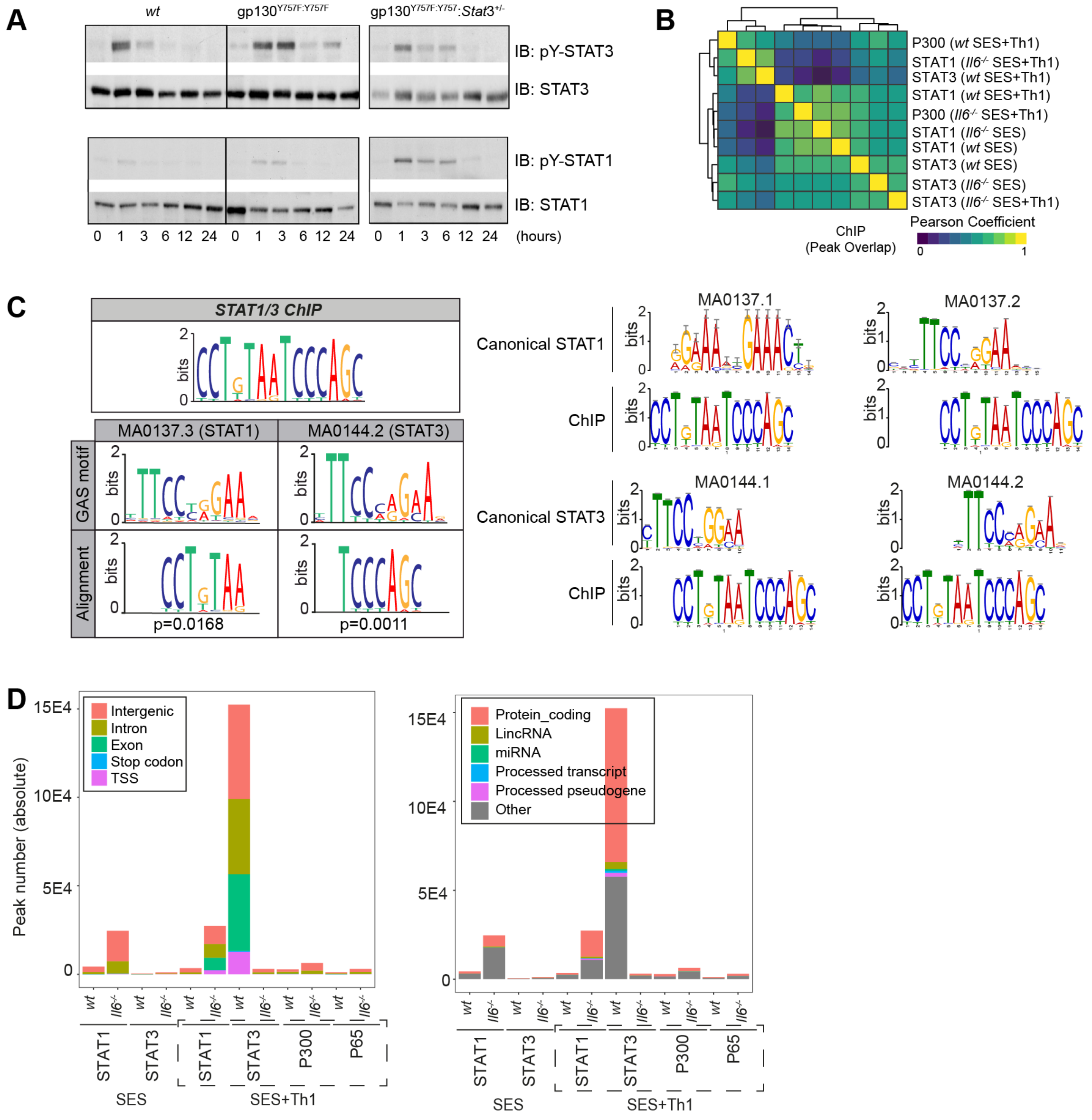

**Supplemental Figure-4**

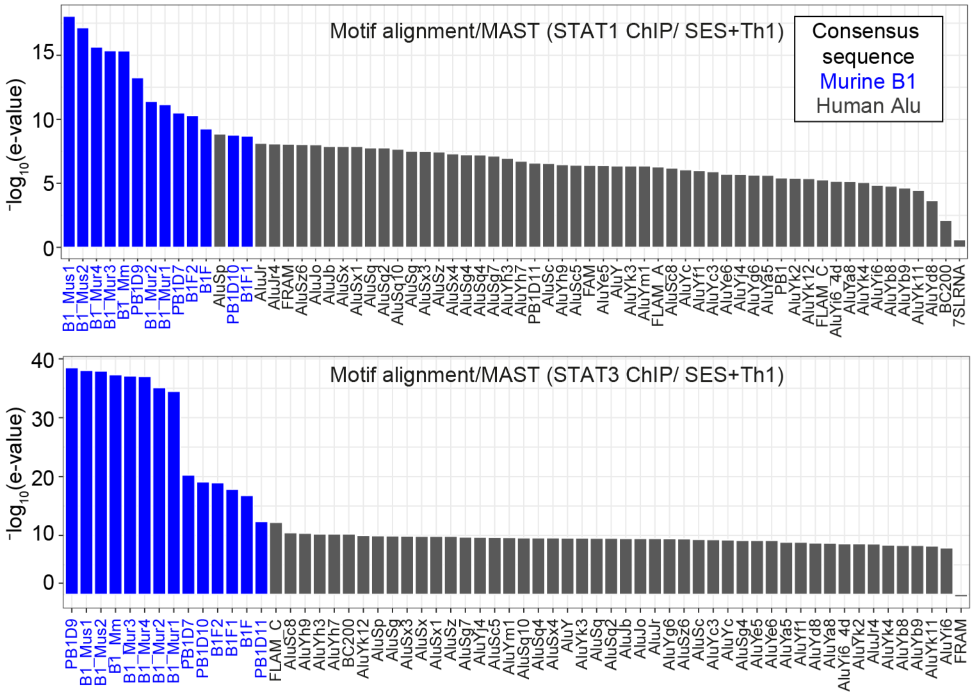

### Supplemental Figure-5

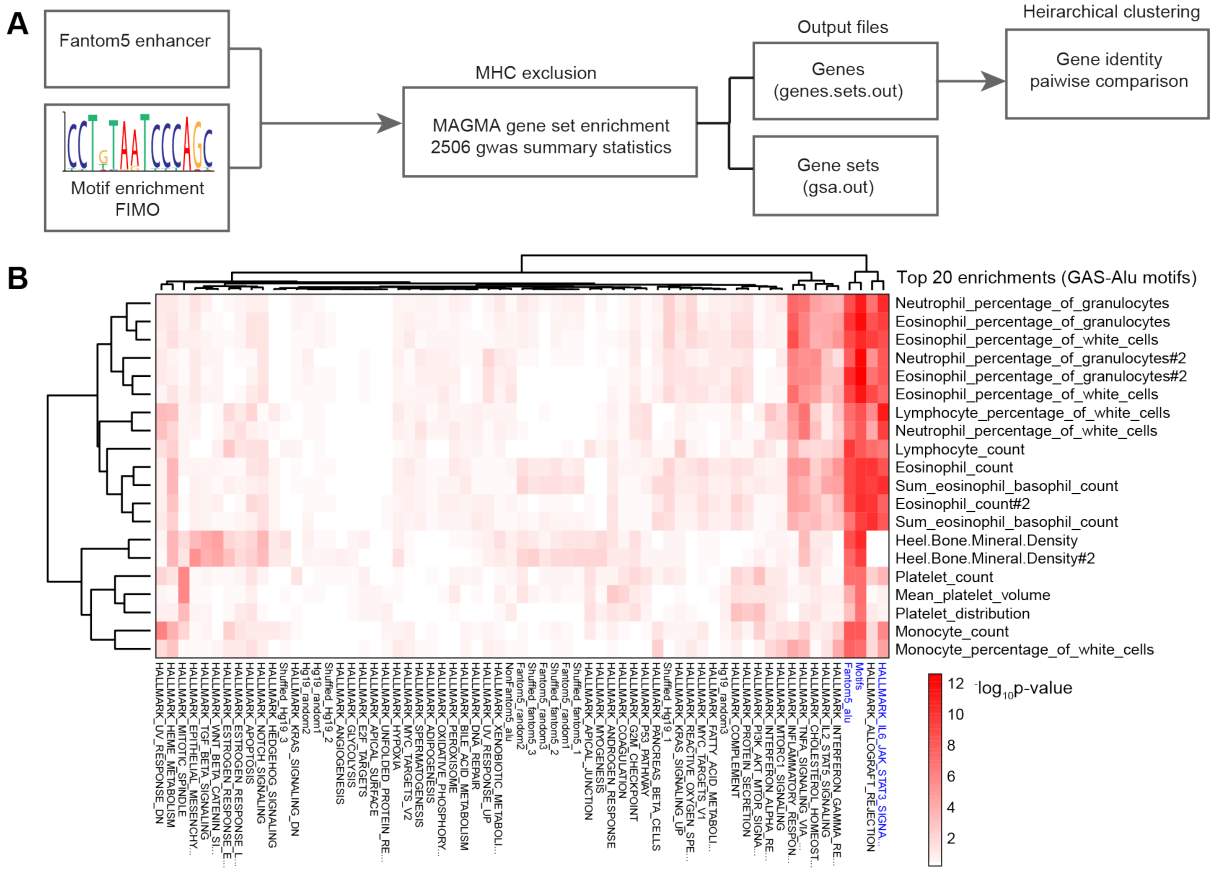
